# Discovering flexible codes for prediction across timescales in the retina

**DOI:** 10.1101/2025.09.19.677348

**Authors:** Kyle Bojanek, Baptiste Lefebvre, Jared Salisbury, Olivier Marre, Stephanie E. Palmer

## Abstract

The retina must encode visual information in a way that supports fast, predictive behavior despite significant processing delays. How this encoding adapts in an ever-changing world, when the temporal statistics of visual input shift, remains an open question. Here we record from populations of retinal ganglion cells in the axolotl as they respond to a stochastic moving bar stimulus across five different temporal correlation scales. Using the information bottleneck (IB) framework, and treating the prediction horizon as a free parameter inferred from the data, we ask what timescale of future motion the retina is optimized to predict under each stimulus condition. We find that the retina adapts its predictive encoding to the changing stimulus statistics: as the time constant of the stimulus dynamics increases, the inferred prediction horizon lengthens. The population shifts toward encoding more velocity information, and motion anticipation grows, all the while maintaining near-optimal prediction efficiency. Population surprise, quantified through a Boltzmann machine model of the retinal response distribution, tracks stimulus surprise under the inferred optimal compression. This connects the retina’s reversal response to efficient predictive encoding. These results show that retinal population codes flexibly adjust their predictive timescale to the temporal structure of their inputs. More broadly, they demonstrate that the IB framework can be used to discover, not just test for, computational objectives in sensory populations.

## INTRODUCTION

Efficient coding, the theory that a population of sensory neurons should maximize mutual information between the response *R* and the stimulus *S* subject to constraints on neural firing rates (Fig. 1), is one of the most influential normative theories of sensory processing [4, 6]. In linear systems with Gaussian statistics, information is determined by the correlation between stimulus and response, but for more complex stimuli and responses, mutual information provides a general measure [17]. Mutual information, *I*(*R*; *S*) = *H*(*R*) *H* − (*R*|*S*), quantifies how much uncertainty about the response *R* is reduced by knowing the stimulus *S*. Maximizing *I*(*R*; *S*) can be achieved either by increasing response entropy (capacity), *H*(*R*), or reducing noise entropy, *H*(*R*|*S*). Maximizing mutual information includes many strategies based on both maximizing population entropy *H*(*R*) (including stimulus decorrelation [1, 3, 6] and high entropy response to input features [9]) and minimizing conditional entropy *H*(*R*|*S*) by decreasing response noise [7, 54]. Information maximization has effectively explained aspects of retinal ganglion cell responses, including center-surround receptive field structure [2], contrast sensitivity functions [3], retinal mosaic geometry [20, 35, 37, 55], response nonlinearities [52], response statistics [5], and cell-type diversity [50].

**FIG. 1.**
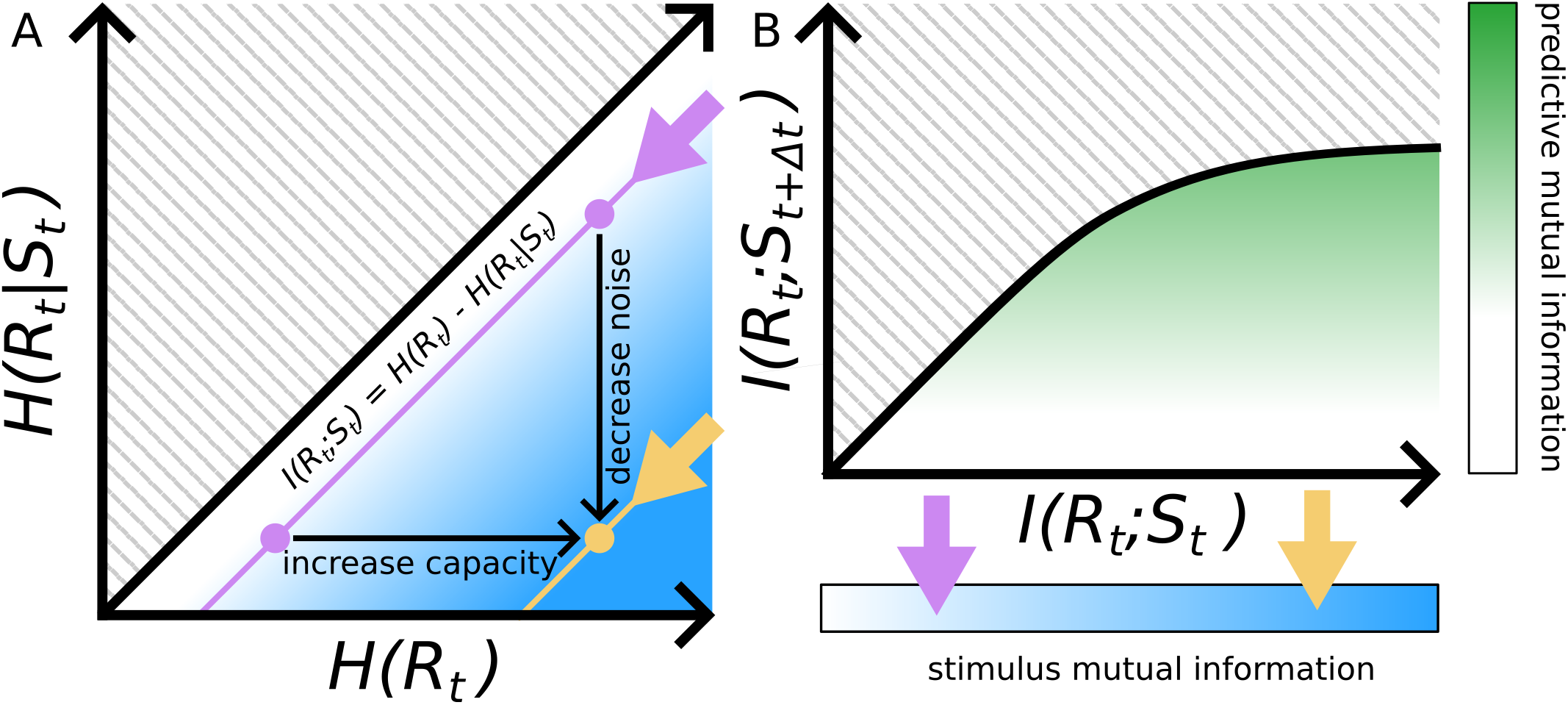
Efficient coding and IB evaluate complementary aspects of sensory coding. **A)** Mutual information, *I*(*R*_*t*_; *S*_*t*_), between stimulus, *S*_*t*_, and response, *R*_*t*_, decomposes as *H*(*R*_*t*_) − *H*(*R*_*t*_|*S*_*t*_). Efficient coding strategies can improve *I*(*R*_*t*_; |*S*_*t*_) either by increasing *H*(*R*_*t*_), for example through stimulus decorrelation or diversifying response patterns, or by decreasing *H*(*R*_*t*_ *S*_*t*_) through noise reduction. Pink dots illustrate different codes with the same *I*(*R*_*t*_; *S*_*t*_), showing that the same information can be achieved at different points along this decomposition. **B)** In the IB framework, a relevance variable (here, the future stimulus *S*_*t*+∆*t*_), is introduced, and the goal is to maximize *I*(*R*_*t*_; *S*_*t*+∆*t*_) for a given amount of information, *I*(*R*_*t*_; *S*_*t*_), retained about the past. The IB bound defines the maximum achievable predictive information at each compression level. Pink dots show codes different codes with the same amount of information, and how they can be improved, arriving at the yellow dot. When evaluating neural codes for optimality, efficient coding principles can increase the position of the neural code on the IB plane.

Classic efficient coding maximizes mutual information between response and the stimulus generally, and does not distinguish which stimulus features are most important for driving behavior. This distinction is especially important in the retina, where slow integration time constants in rods and cones [8, 54] have the potential to cause behaviorally detrimental visual delays. Features that allow the system to anticipate the future are therefore particularly important. This idea can be formalized using the information bottleneck (IB) framework, a technique from information theory, where information *about* some desired relevance variable is maximized, subject to a constraint on the total amount of information available [13, 14, 65] (Fig. 1). For anticipation of stimuli, the relevance variable is taken to be the future stimulus state *S*_*t*+∆*t*_ (see Methods). This choice of relevance variable naturally leads to delay compensation. Analysis of retinal ganglion cell responses using the IB framework [41, 51] reveals that small populations of retinal ganglion cells sit near an upper bound on the maximum possible information between the future stimulus and the population response given their measured amount of information about the past.

Successful applications of efficient coding and IB suggests that both may capture important aspects of retina population activity. For versions of efficient coding that do not constrain what features should be encoded, the two analyses can be complementary: efficient coding constrains *how* information is represented, while the IB framework determines *what* information is represented. In the simplest terms, efficient coding asks whether the available channel capacity is used well (e.g. are spikes wasted on noise or redundancy?), while IB asks whether the bits that are transmitted are the right bits for a given task. For prediction, the right bits are those that carry information about the future stimulus. The two frameworks can be evaluated on the same data: a population that sits near the IB bound is encoding the right features, and a population with high efficiency (low noise entropy, decorrelated responses) is using its capacity well. A code can be IB-optimal but wasteful of channel capacity, or highly efficient but encoding the wrong features. Here we evaluate both. In the predictive setting, the question of, “What is encoded?” depends on the choice of timescale, ∆*t*_IB_, which sets how far into the future the relevance variable extends. Previous work fixed ∆*t*_IB_, but here we infer 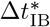 from the population response across different stimulus conditions. This establishes a novel way to examine how the retinal encoding adapts to changing stimulus statistics.

The Deterministic Information Bottleneck formalizes another notion of optimality [64]. Consider two neural populations that carry the same mutual information *I*(*S*; *R*) between stimulus and response. One code achieves this with sparse spiking, and thus low response entropy *H*(*R*), while the other uses high firing rates, producing high *H*(*R*). Since *I*(*S*; *R*) = *H*(*R*) − *H*(*R*|*S*), the high-entropy population must also have high conditional entropy *H*(*R*|*S*): more of its response variability does not encode stimulus information. From a classic IB perspective, the two codes are equivalent, but the Deterministic IB favors the lower-entropy solution, which uses fewer spikes to convey the same information. This preference is especially relevant for the retina, where responses are dominated by silence. Here, we evaluate retinal population responses along both axes: proximity to the IB bound (encoding the right features) and efficiency of the encoding (using capacity well) (Fig. 1). This analysis is done across a range of stimulus statistics to examine if the 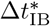 that best describes the retina population response adapts to changing stimulus statistics. For retinal populations, the optimal 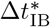 increases as a function of the stimulus time constant *τ*; across all stimulus conditions the population decoding performance is well described by IB theory.

## RESULTS

### What is the appropriate relevance variable?

Teasing apart how stimulus features affect predictive encoding requires a rich, yet tractable input motion model. A moving bar stimulus whose dynamics are generated by a stochastically driven damped harmonic oscillator (Fig. 2a), is a physically motivated choice of dynamical system that has exactly solvable correlation functions [48] while still displaying rich behaviors useful for probing the retina, like motion reversals (Fig. 2a). In addition to the correlation functions being exactly solvable, the IB problem for future information can also been solved exactly [51] across all choices of stimulus parameters [56]. Additionally, the stimulus is Markovian and therefore optimal estimation of the future state *S*_*t*+∆*t*_ = (*x*_*t*+∆*t*_, *v*_*t*+∆*t*_) depends only on it’s present position *x*_*t*_ and velocity *v*_*t*_, which constitute the stimulus state space *S*_*t*_ = (*x*_*t*_, *v*_*t*_) (Fig. 2b). Therefore, the information bottleneck encodings *T* are noisy (*ξ* = 𝒩(0, *I*)) linear combinations of present stimulus position and velocity, *T* = *AS*_*t*_ + *ξ* [15]. The choice of encoder map *A* can be understood in terms of the performance of an optimal decoder of the present state *P* (*S*_*t*_|*T*) and how well it decodes the future state *P* (*S*_*t*+∆*t*_|*T*), all of which are analytically tractable for our choice of stimulus (see Methods). It is therefore possible to use the performance of the optimal decoder given *T* and the resulting distribution of errors to understand different possible encodings *T*. Consider two different encoding schemes that are equally informative about the stimulus at time ∆*t* = 0 (Fig. 2c, top and bottom). At equal time both carry the same amount of information about the stimulus, however one encoder results in anti correlated errors (Fig. 2c top) in decoding position and velocity, while the other has correlated errors (Fig. 2c, bottom). While the two have the same initial information about the stimulus (∆*t* = 0), the encoder that results in correlated errors loses information much faster as ∆*t* increases (Fig. 2c).

**FIG. 2.**
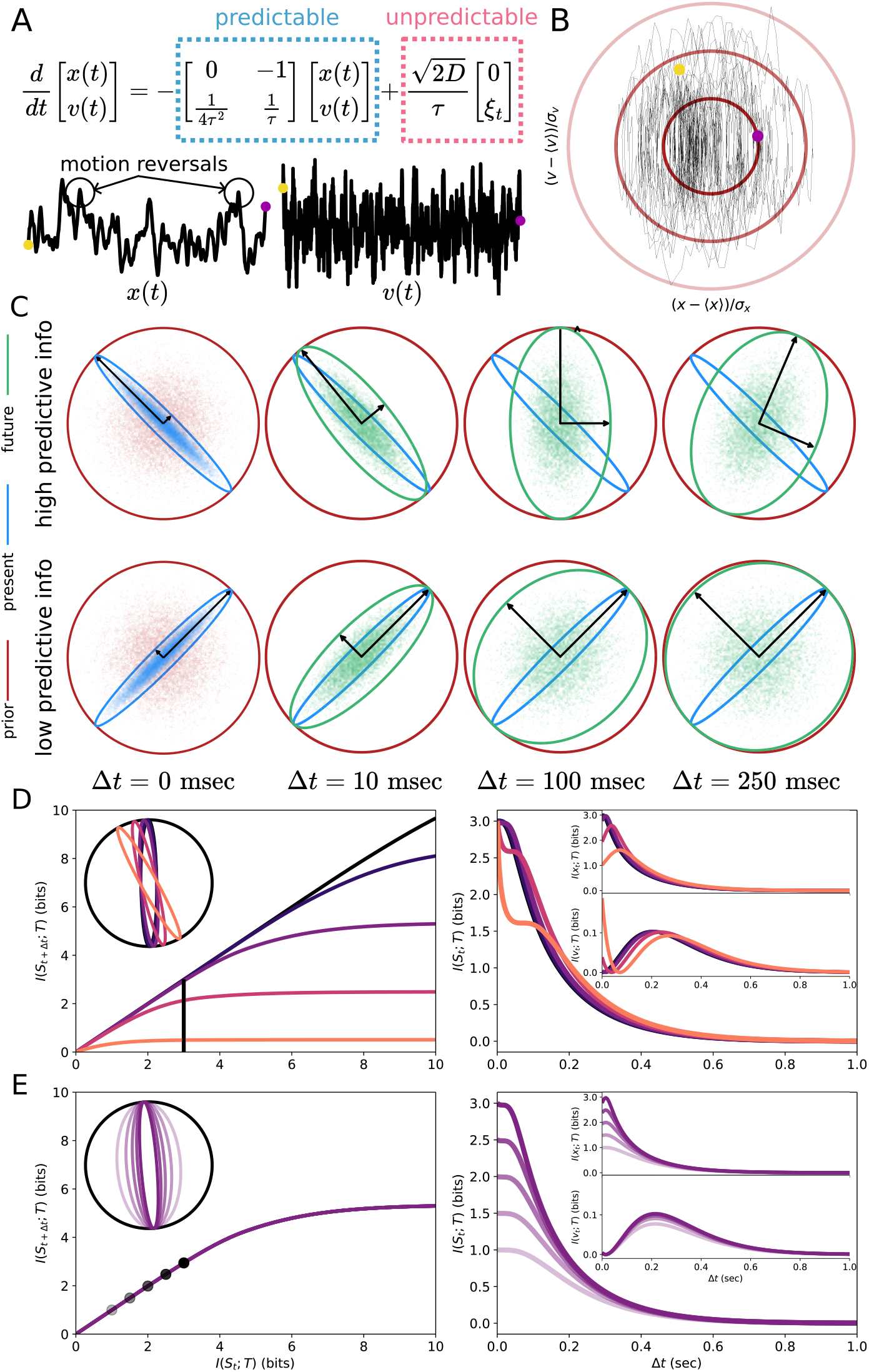
Connection between motion anticipation and the predictive IB. **A**) A stochastically driven critically damped harmonic oscillator controls the position of a white bar on a black screen. The dynamics include both a predictable component (oscillator) and an unpredictable component (white noise drive). Example position and velocity traces are shown for *τ* = 100 ms **B**) along with the corresponding phase space. Concentric ellipses in phase space show level sets for 1, 2 and 3 *σ*. **C**) Schematic illustrating the effect of different measurements of position and velocity information. Three-standard-deviation ellipses represent prior, present measurement, and future uncertainty across a range of times. These ellipses represent the distributions of errors made by an optimal decoder. Samples of the errors are shown as scatter points. **D**) The effect of changing ∆*t* (from dark to light evenly log-spaced from 3ms to 316 ms) in the IB problem is shown for the information bottleneck plane, and the amount of total, position, and velocity information. **E**) The same is shown for fixed ∆*t* and changing the information about the present *I*(*S*_*t*_; *T*).

Applying IB to the stimulus prediction problem requires choosing how far out in the future to predict. Sachdeva et al. [56] demonstrated that when the relevance variable is the future stimulus state (position and velocity) at time *t*+∆*t*_IB_, the optimal encoding smoothly transitions from encoding position information (small ∆*t*_IB_) to incorporating increasing amounts of velocity information (large ∆*t*_IB_) (Fig. 2d inset). This creates a family of optimal encodings as ∆*t*_IB_ varies from near zero to infinity. Additionally, as ∆*t*_IB_ increases, the total amount of information defining the IB upper bound decreases, as noise degrades the amount that can be know about the future as ∆*t*_IB_ increases (Fig. 2d). In the high-compression regime, the two-dimensional state space is projected down to one dimension, and the optimal encoding is a linear combination of position and velocity with some Gaussian noise injected *ξ*. When ∆*t*_IB_ is allowed to vary, IB optimal encodings span a substantial fraction of all possible linear combinations of position and velocity encoding. The space of potentially optimal encodings grows even larger when alternative relevance variables are considered, such as future velocity alone rather than the full future state. In the extreme, one could even construct an artificial relevance variable that makes any given retinal encoding match the IB-optimal one by choosing the relevance variable to match exactly what the retina encodes. This emphasizes how the IB optimum is often matchable to neural data for a careful choice of relevance variable, revealing what computations the brain may be optimized for.

To address the large number of potentially optimal encoding maps, our analysis of the retina population response begins by constructing the entire set of predictive IB encodings, which vary as a function of ∆*t*_IB_. Then, depending on which ∆*t*_IB_ best describes the population response, that 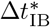 defines the relevance variable 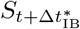 (see Methods). This extends IB from a formalism for studying the optimal encoding of information, to a tool that can be used to learn something about the population response. The 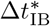 that best describes the population response suggests that particular value is what the retina has evolved to predict. It would be reasonable to conclude that two different retina populations that were well described by two different values of 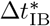 may have different output functions. The feature that the retina is encoding best in response to a stimulus may be considered an answer to the question, “What is the retina computing?”

### Motion anticipation and IB

IB is an information theoretic tool, however for the choice of stimulus we make here, it has a straight-forward interpretation in terms of motion anticipation. Motion anticipation in the retina has previously been probed with deterministic moving bar stimuli, traveling at constant velocity [10, 67, 68] and constant speed with sudden reversals [16, 60]. The effect can be extended to a more ethologically relevant, stochastic moving bar stimulus [51]. For the deterministic moving bar stimulus, motion anticipation is defined as the location in space where the retina response is peaked relative to the leading edge of the bar stimulus itself. For a stochastic stimulus, we define motion anticipation as follows: given the population response at time *t*, we compute the information between a compressed stimulus representation and the actual bar position at time *t* +∆*t*, across a range of ∆*t*. If the peak of information occurs at positive ∆*t*, meaning the population response at time *t* is most informative about a future stimulus position, the population response is anticipatory. The amount of anticipation is the time shift of this peak relative to zero.

The solution of the IB problem for the stimulus shows that as the relevance variable defined by 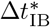 moves further into the future, the optimal encoding shifts the time of peak position information forward in time; the compression leads the stimulus by a larger amount of time. The data processing inequality [17] (in this context, it is impossible to know more about the future than the present) implies the total information about the stimulus must monotonically decrease as time marches forward (Fig. 2d). However, information about position alone can increase as a function of the time offset (Fig. 2d). This increase is motion anticipation. Similarly, information about velocity under optimal IB compression is not monotonically decreasing (Fig. 2d). Thus, motion anticipation [10], a classic effect observed in the retina, appears here as the IB optimal encoding of a stochastic stimulus. It is also notable that this motion anticipation comes at a cost: at fixed levels of stimulus information, the anticipatory response comes from trading some position information for velocity information. Thus, as the timescale of anticipation increases, the initial position decoding performance must also decrease (Fig. 2d). The total amount of anticipation in the information bottleneck solution can be solved for analytically (see Methods).

It is also possible to examine the effect of changing the total amount of information the IB compression has about the stimulus for fixed ∆*t*_IB_. In this case (in the limit of high compression), the relative amounts of position and velocity information are fixed, and so the time of peak position information is stable (Fig. 2e).

### Evaluating the optimality of retinal response to stimuli with different time constants

With this framework in hand, a population of 53 axolotl retinal ganglion cells is analyzed, as it responds to a stochastically driven critically damped harmonic oscillator with 5 different choices of time constant *τ* (25 ms, 50 ms, 100 ms, 200 ms, 400 ms) (Fig. 3a,b). Response are assessed for optimality under both efficient coding and predictive IB.

**FIG. 3.**
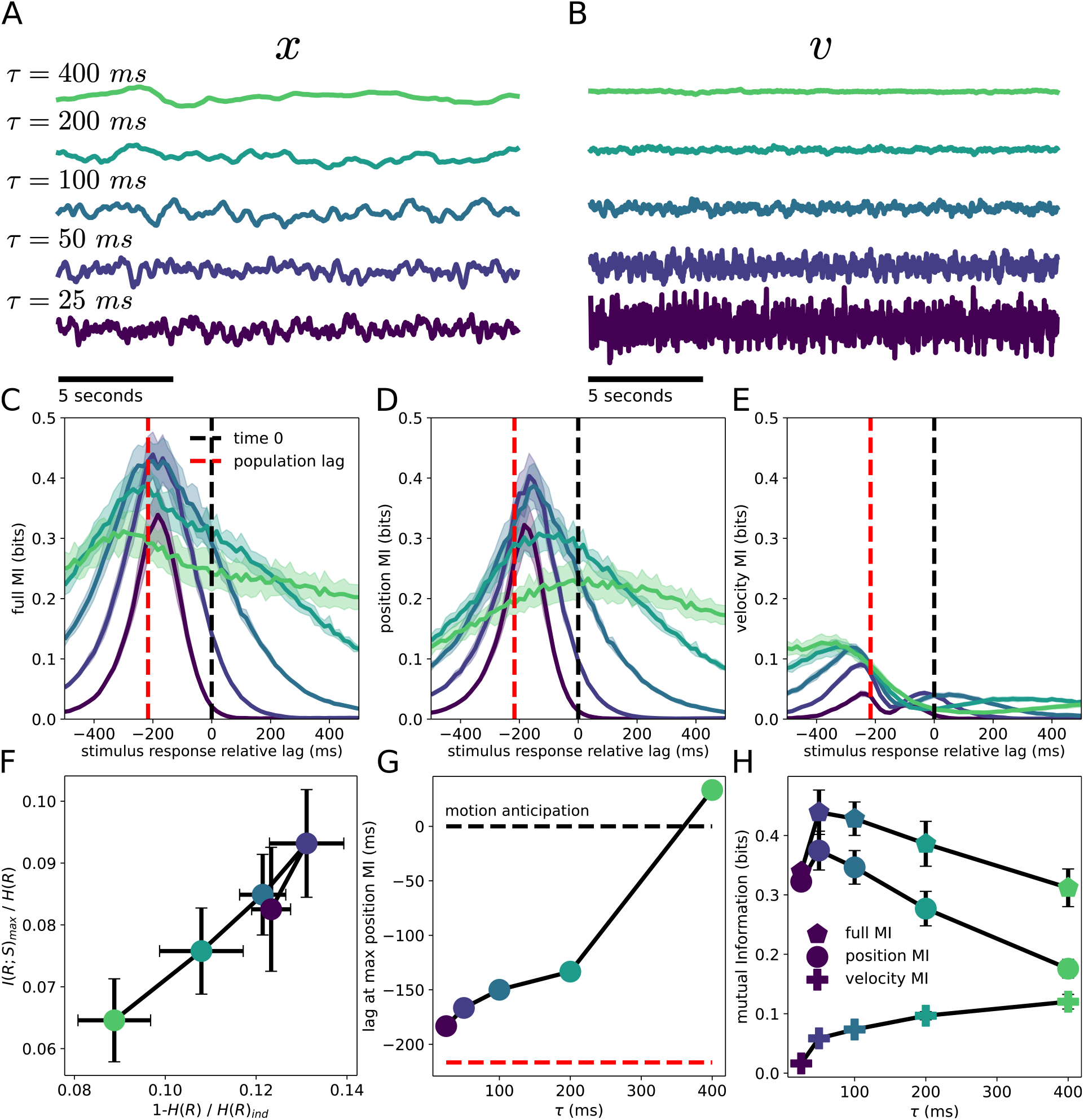
Relative priority of position and velocity information changes with stimulus time constant. **A)** Example bar position traces for different time constants. Noise is adjusted to keep bar position variance constant. **B)** Corresponding bar velocity traces. With position variance fixed, velocity variance decreases as the time constant increases. **C-E)** Mutual information between one bin (16.667 ms) of neural activity and (**C**) the full stimulus, (**D**) position, and (**E**) velocity. Average response delay remains consistent for total information (**C**). As *τ* increases, the time of maximum position information shifts forward (**D**), reflecting increased velocity information (**E**). Red dotted line: time of peak mutual information between response and stimulus averaged across conditions. Black dotted line marks time 0. **F)** The two coding-efficiency measures plotted against each other. The y-axis, 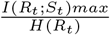 quantifies noise in the population response; the x-axis 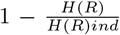 measures stimulus decorrelation. The least noisy condition is also the most strongly correlated. **G)** Time to peak position information as a function of *τ*. Because responses carry velocity information in all conditions, peaks are consistently later than total information peaks. The peak shifts further forward with increasing time constant, extending into the future for the *τ* = 400 ms condition. **H)** Decomposition of total mutual information into position and velocity contributions as a function of time constant. Increasing time constant emphasizes velocity information, enabling greater motion anticipation. Error bars are given by standard deviations over folds

The mutual information is measured between the population stimulus at a give time, *t*, and the position and velocity of the stimulus across a range of relative times in the past or future, *I*(*S*_*t*_; *R*_*t*+*k*_). Mutual information quantities are estimated using the performance of position and velocity decoders as a variational lower-bound on the true conditional entropy *H*(*S*_*t*_|*R*_*t*+*k*_) (see Methods). Reported information quantities are the average performance over test data; error bars are standard deviations over multiple folds of testing data. All stimulus conditions show a similar response delay, measured as the time of peak mutual information between the stimulus and response (Fig. 3c). While mutual information between the population response and the full stimulus (position *and* velocity) has a somewhat consistent time of peak information (Fig. 3c), there is a much less consistent peak in position-only information (Fig. 3d). As the time constant of the stimulus grows larger, the time of peak position information moves farther and farther forward in time (Fig. 3d). For the slowest time constant stimulus (*τ* = 400 ms), the population response becomes anticipatory (Fig. 3g). That is, the time at which the population best decodes the bar position occurs in the future. This happens because the neural population response carries more and more information about the velocity of the stimulus, allowing for extrapolation (Fig. 3e).

This increase in the time of peak position decoding performance as the time constant of the moving bar increases is consistent with 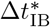 increasing in the predictive IB problem. As the time constant of the stimulus grows, more information is carried about velocity, and less about position (Fig. 3h). The application of IB makes clear how the population response of the retina changes as a function of stimulus parameters. A different measure of optimality needs to be considered to evaluate the population response in terms of informational efficiency.

First, the amount of noise in the system can be quantified 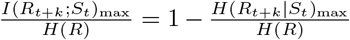. This measure will be 1 for a system with no noise, and 0 for a system with only noisy responses. Maximizing this measure corresponds to maximizing information between the stimulus and response by minimizing the conditional entropy, *H*(*R*|*S*). In the retinal data, the population response has a maximum in efficiency for *τ* = 50 ms (Fig. 3f). Second, efficient coding in terms of stimulus decorrelation is measured by one minus the ratio of actual system entropy *H*(*R*) to the entropy of a rate-matched independent population *H*(*R*_ind_). This measure, 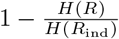will be 0 for a fully independent population, and increase as correlations grow stronger. This condition normalized multi-information [58] measure is appropriate for comparison across stimulus conditions because stimuli were variance-matched across *τ*, meaning that the level of correlation between neurons should be a result of retinal response properties, rather than stimulus properties. Under this measure, a maximum is observed for *τ* = 50 ms (Fig. 3f). Interestingly, while the noise reduction measure is at a maximum, the population response is the most strongly correlated 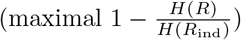.

It is important to consider the relative magnitude of these competing effects. In terms of capacity lost due to correlations between neurons, in order of increasing *τ*, a reduction in entropy of 0.58±0.06 bits, 0.72 ± 0.03 bits, 0.72 ±0.03 bits, 0.63± 0.05 bits, and 0.48± 0.06 bits is observed. Measuring the information lost due to noise, *H*(*R*|*S*), we observe 3.8 ± 0.29 bits, 4.35 ± 0.14 bits, 4.80± 0.18 bits, 4.80± 0.46 bits, and 4.66± 0.29 bits (stimulus conditions in the same order). While decorrelation of the stimulus has the potential to improve information between stimulus and response, it appears that for the level of noise in the retina, much larger gains are possible with noise suppression.

The *τ* = 50 ms stimulus condition achieves maximal mutual information between stimulus and response (Fig. 3c), while still having the second lowest entropy of all stimulus conditions, and the least-independent population response. The superior performance over other stimulus conditions is a result of less noise in the population during this input drive. This suggests an interesting response regime for the retina in which correlations between retinal ganglion cells are (at the very least) not harmful, if not beneficial, for stimulus encoding. While this analysis is carried out for single time points in the stimulus and response, retinal ganglion cells populations carry information about the stimulus across time, which can have important implications for retinal coding [44]. The same analysis was performed decoding from multiple time bins, up to 500 ms in duration, and similar effects are observed (data not shown).

Because the measured relative amounts of position and velocity information in the retina response change across stimulus conditions (Fig. 3c-e), there is no one choice ∆*t*_IB_ in the *S*_*t*+∆*t*_ relevance variable that the retina response would be optimal for. Therefore to assess IB optimality we infer a value 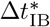 to define the relevance variable 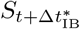 that best describes the retina population response (see Methods). For this optimization we take the time in the future that the retina has maximal information about the present of the stimulus as a processing lag, and consider IB optimality relative to some lag *k*, similar to [41]. Placing the retina population response relative to the upper bound on future information set by the information bottleneck for the inferred 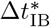, we find that in all cases the retina population response is near the bound (Fig. 4a). Additionally, since all linear measurements of the stimulus position and velocity still give information about the future stimulus state, even if sub-optimally, there is therefore a lower bound on the amount of predictive information [43]. In all cases the retina population response is far from the lower bound on predictive information.

**FIG. 4.**
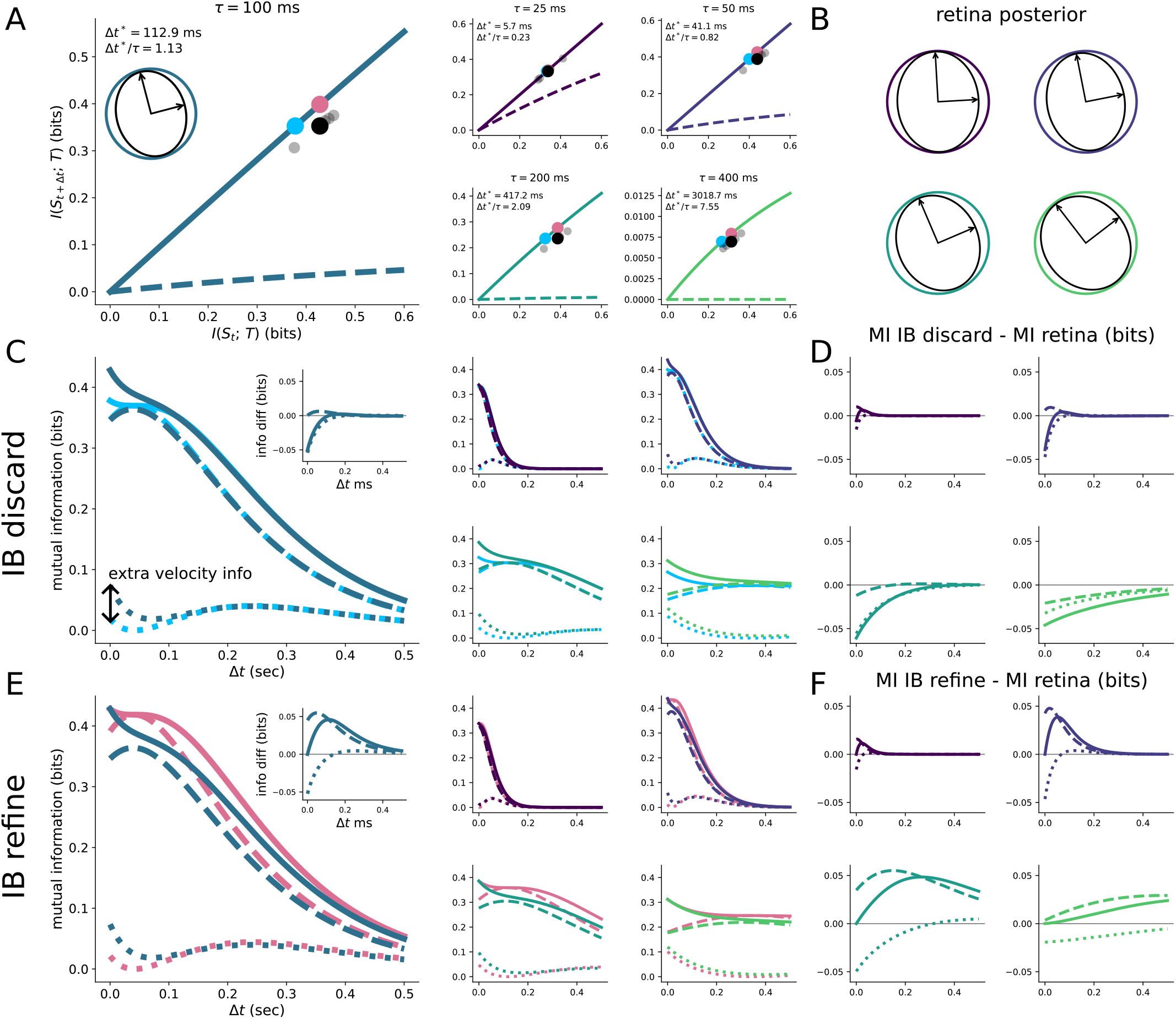
IB encodings accurately describe population responses. **A)** Position of retina population response relative to IB upper bound is shown across stimulus conditions. Black dot is average over test folds for estimating information; gray dots are for individual folds, the blue dot identifies the IB discard solution, and the red dot identifies the IB refine solution. Solid line shows IB (upper bound) solution, the dashed line shows the lower bound on predictive information. **B)** Implied posterior from decoding analysis is shown for each stimulus condition. **C)** Total, position and velocity information are shown as a function of time for both the retina population response, and the an IB optimal solution that achieves equal performance using less information (IB discard) across stimulus conditions. **D)** The difference between the stimulus and IB discard optimal information are shown for total, position and velocity information as a function of time across all stimulus conditions. **E)** Total, position and velocity information are shown as a function of time for both the retina population response, and the an IB optimal solution that achieves superior performance using the same amount of information (IB refine) across stimulus conditions. **F)** The difference between the stimulus and IB refine optimal information are shown for total, position and velocity information as a function of time across all stimulus conditions.

The value of 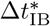 that best describes the population increases as a function of *τ*, consistent with the increasing amounts of velocity information (Fig. 3h). This is true in absolute terms 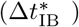and in relative terms 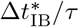 (Fig. 4a). The effect of increasing velocity information across stimulus conditions can also be seen in the posterior of the decoder *P* (*S*_*t*_|*R*_*t*+*k*_) (Fig. 4b). To understand the effect of information bottleneck sub-optimality, we consider two hypothetical IB optimal encoders, “IB discard” and “IB refine.” “IB discard” (Fig. 4c,d) moves to the IB optimal bound by removing information about the stimulus to achieve the same level of performance as the retina population response. Across stimulus conditions the retina response loses position information in a manner very similar to the “IB discard” solution, but the retina also carries a small amount of velocity information beyond the minimum necessary to be IB optimal (Fig. 4c,d). “IB refine” moves to the IB optimal bound by reshaping the stimulus representation to have the same amount of information about the stimulus while achieving the maximum amount of predictive information (Fig. 4e,f).

When an extra degree of freedom is allowed for how far out into the future the population of retinal ganglion cells is predicting, we can observe near optimal IB encoding across a range of stimulus conditions. A fixed choice of IB relevance variable would not have described the retina response to all stimulus conditions well. This suggests a novel approach to studying adaptation effects on encoding at a high level.

### Implications of efficient coding with surprise

One prediction made by efficient coding theory, particularly predictive coding [23, 33, 53], is that neurons encode the novelty of their input. The dynamic stimulus analyzed here requires a careful treatment to quantify surprise in a probabilistic sense. While many notions of statistical surprise exist [46], a common and well-motivated choice is the negative log-probability of the input-essentially how ‘rare’ an input is. This conception of stimulus novelty was found to describe the firing patterns of individual retinal ganglion cells in response to a full field flash stimulus designed to evoke the so-called Omitted Stimulus Response [19, 59]. Estimates of the population response log-probability are used to extend this quantitative analysis explored with single cells as in [19] to the entire population.

As the retina highly compresses incoming stimuli, we must consider not the novelty of the stimulus, but rather the novelty of the compressed stimulus. This means that the surprise of the population response, estimated independently from a Boltzmann machine model, should track the surprise of the IB-compressed stimulus. This is the connection we test below.

The negative log-probability of the retina population response is computed and compared to the negative log-probability of both the stimulus *P* (*S*_*t*_) and the compression of the stimulus found from our decoding analysis *P* (*T*|*S*_*t*_), putting both input and output ‘surprise’ in direct comparison. As the lag for the comparison and the compression map were set based on the decoding analysis, there are no free-parameters for comparing the stimulus and compression surprise to the population response surprise. We perform a head-to-head comparison of the stimulus and response surprise (Fig. 5a).

**FIG. 5.**
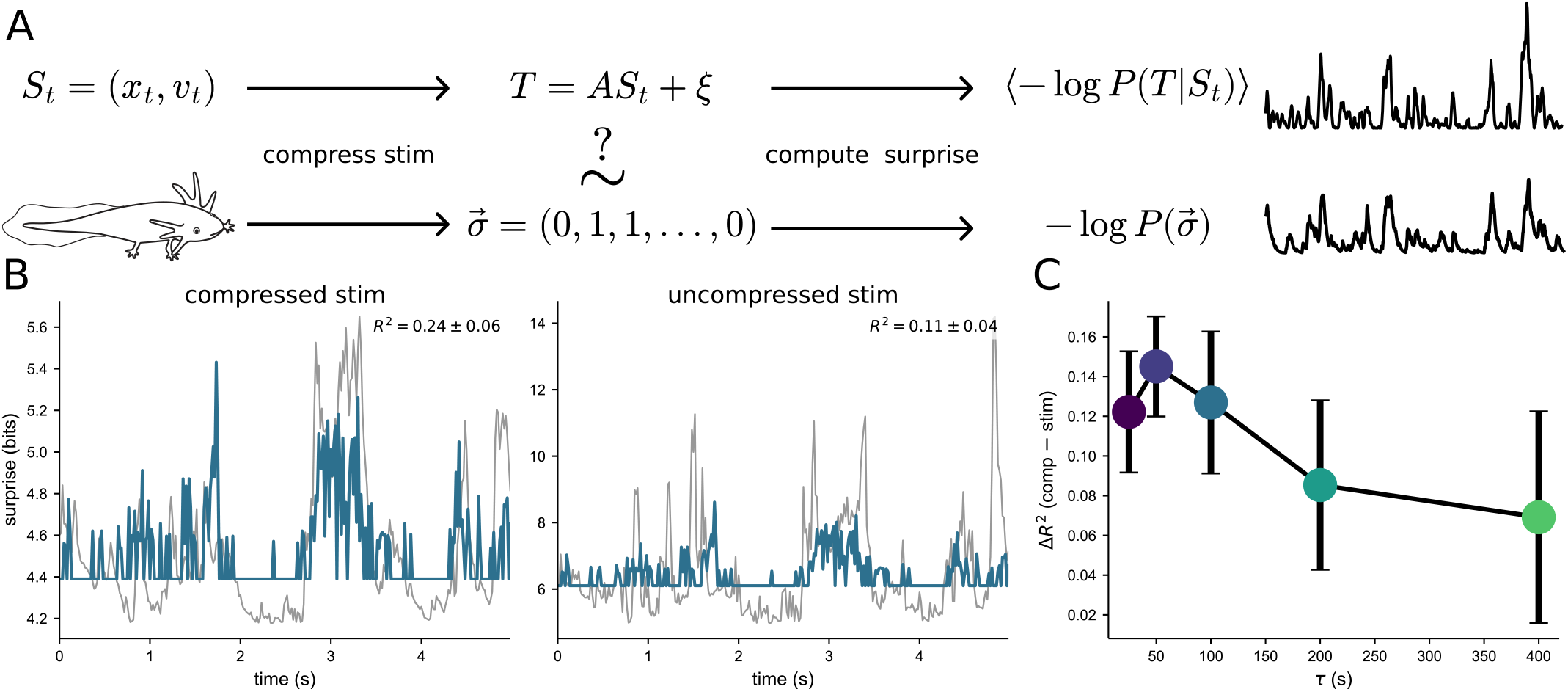
Surprise encoding and the retina reversal response. **A)** Schematic of the procedure for comparing stimulus surprise with response surprise. **B)** Relationship between retina response surprise (blue), and compression surprise (left, gray) and stimulus surprise (right, gray). *R*^2^ means are given by average over stimulus trials along with standard deviation over trials. **C)** Difference between compressed *R*^2^ and uncompressed *R*^2^ (additional variance explained) is shown for all stimulus conditions.

To estimate the probability of each population response pattern, and therefore population surprise, we need a model of the full joint distribution *P* (*R*). The decoder gives us *P* (*S*|*R*) but not *P* (*R*), itself. We fit a Restricted Boltzmann Machine (RBM) to the population spiking patterns, a flexible energy-based model that has been shown to accurately capture the statistics of retinal population activity, including higher-order correlations beyond pairwise interactions [24]. The RBM provides an estimate of −log *P* (*R*) for each observed response pattern, which we take as a measure of population surprise. Across all stimulus conditions tested, the correlation between the stimulus compression and response log-probabilities is significantly stronger than that between the uncompressed stimuli and the response log probabilities (Fig. 5b,c). indicating that a surprising input leads to an especially rare and surprising response that could potentially be used to signal surprise to downstream circuits. Traces shown are for the *τ* = 100 ms stimulus condition (for other conditions see Methods). This ability to decode stimulus surprise directly from population surprise demonstrates that the efficient coding theory prediction that rare responses should occur with rare stimuli is borne out across stimuli conditions, but only for the compressed stimuli.

This observation suggests a tight a connection with the retina reversal response [60], a phenomenon observed in retinal ganglion cells where the sudden reversal of a moving bar evokes a large and synchronous volley of spikes. Since the statistics of the stimulus are known, we can compute the stimulus reversal probability as a function of position and velocity. Compressions carrying predictive information are more informative about reversal probabilities than non predictive compressions (see Methods).

## DISCUSSION

We have shown three main results connecting different notions of efficient coding to retinal responses to a suite of moving bar stimuli. First, by treating the prediction horizon ∆*t*_IB_ as a free parameter, the IB framework becomes a discovery tool. We infer from the data what timescale of prediction best describes the retinal population code, rather than testing optimality at a single fixed (guessed) timescale. Second, as stimulus statistics change, specifically as the time constant of the driving dynamics increases, the inferred prediction horizon 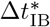 increases in parallel, and the population code remains near the IB bound across all conditions. This is evidence that the retina changes its predictive encoding to the temporal structure of its inputs. Third, population surprise, estimated from a Boltzmann machine model, tracks stimulus surprise under the inferred IB-optimal compression, linking the framework to the retina’s reversal response and to classical predictions of efficient coding theory.

An information theoretic interpretation of classic retinal ganglion cell response features (motion anticipation and reversal response) using IB and efficient coding arguments shows how both of these notions of “optimal” can be simultaneously evaluated in data. When performing this kind of analysis in the context of prediction in time, it is important to consider laws of mutual information such as the data processing inequality. A time offset can be used to account for delays in the retina response, but this offset must be chosen in such a way that respects causality. If an offset is chosen too far in the past, apparent (but, of course, spurious) violations of these laws can appear. Under choices like this, mutual information between the stimulus and response can increase from the time chosen as the offset, which would be divination.

An important question is how, mechanistically, these optimal and efficient computations are carried out by populations of retinal ganglion cells. Previous work has highlighted the role of amacrine cells in motion anticipation [34], and other studies suggest that gap junctions may contribute to predictive encoding of information [41], with anticipatory effects also observed in bipolar cells, likely mediated through interactions with amacrine cells [41]. A common theme among these possible mechanisms is inhibition [22], either feedback or feedforward. Theoretical analysis has made specific predictions about speed dependence of motion anticipation in feedback and feedforward inhibition [22]. Careful measurement of motion anticipation effects across a range of stimuli, similar to what is reported here, could be useful in determining whether feedback or feedforward inhibition accounts for motion anticipation effects observed in populations of retinal ganglion cells.

Another open mechanistic question is how single cell biophysics may contribute to predictive processing observed in the retina. This has been explored in phenomenological models [10], but has not been linked explicitly to single cell mechanisms. Spike frequency adaptation is a form of negative feedback observed across many different types of neurons [57], which include retinal ganglion cells [49]. Such adaptation effects have been shown to cause phase leads in single neocortical pyramidal neurons injected with sinusoidal current drive [42]. This work was not done in the context of motion anticipation, but the development of phase leads is necessary for motion anticipation as described in prospective coding [11]. For this reason, it may be possible for single retinal ganglion cells to show motion anticipation effects in isolation. While pharmacological knock-outs of circuit mechanisms have effectively removed motion anticipation from retinal ganglion cells [34], suggesting that they are necessary for motion anticipation, it does not show that they are sufficient for motion anticipation. Some amount of predictive capacity may arise from single cell properties. Such effects arising from single cell biophysics would likely play a synergistic role with other circuit based mechanisms.

An important caveat is that the stochastic moving bar stimulus used here, while physically motivated and analytically tractable, is not a naturalistic stimulus for the axolotl. The damped harmonic oscillator captures generic features of inertial motion, e.g. constant velocity epochs, reversals, starts and stops, that are shared with natural motion. Our previous work has shown that such stimuli evoke predictive coding in the retina that generalizes across stimulus classes [21, 32, 39, 70]. Nevertheless, the specific prediction horizons and optimal encodings we infer are properties of the retina’s response to this particular stimulus ensemble. Extending this approach to natural motion stimuli, for which the information bottleneck problem is no longer analytically solvable but may be tractable with recent variational methods, is an important direction for future work. This would test whether the adaptive predictive coding we observe here reflects a general property of retinal circuits. The IB approach outlined here may be useful in other neural systems. By constructing a family of optimal encoding distributions that come from different choices of relevance variables and finding the relevance variable that best describes the system response, one is able to *discover* what the system is optimized for. When analyzing a population response, if there is some parsimonious relevance variable for which the response is optimal, that variable is probably closely linked to the function of the system. While prediction of future stimulus is well-motivated as a family of relevance variables, particularly for the early sensory systems, other choices may be appropriate for different brain areas. Additionally, across species, the retina has a number of specialized functions linked to different cell types; examples include the detection of looming objects [38, 47], and motion detection [18]. More generally, across species there is a variety of more specific, ethologically important behaviors that may require entirely different choices of relevance variables [40].

As a final point, there is exceptional value in using stochastic stimuli in sensory experiments. The stochastic, dynamic, and yet analytically-tractable stimulus used here allows for a set of information theoretic analyses that would have been ill-posed with a deterministic stimulus. In addition, this stochastic stimulus enabled connections to be drawn between the retina reversal response and motion anticipation. Stimuli with rich naturalistic spatial and temporal correlation structure not only evoke the most expressive response states in neural populations, they also allow for the discovery of what matters to the brain and what it may have evolved to encode. Future research may take advantage of the rapid progress being made in generative models of images and videos to extend this probabilistic approach to a more naturalistic setting [29, 36, 63].

## METHODS

### Retina recording

Following methods described in [27], 53 retinal ganglion cells from the isolated retina of an axolotl were recorded using a multielectrode array with 30 µm spacing and 252 channels.

### Retina stimulus

The retinal ganglion cells were presented with a stimulus of a white moving bar against a black screen. The dynamics of the bar were governed by the described stochastically driven critically damped harmonic oscillator. The chosen time scales were 25 ms, 50 ms, 100 ms, 200 ms, and 400 ms. The projection used a DMD system with a refresh rate of 120 Hz. The spikes and stimulus were both binned at 60 Hz. The stimulus conditions were interleaved, and each condition had 30 different trials each lasting 20 seconds.

### Information bottleneck for prediction

The Information bottleneck (IB) [65] is a powerful tool for applying rate-distortion theory [17] to problems in supervised learning. An IB problem assumes an input variable *X*, a noisy compression of the input *T*, and a relevance variable *Y*. The variables satisfy the Markov condition *T* −*X*−*Y*, that is, *T* is conditionally independent of *Y* given *X*, meaning all information that *T* has about *Y* flows through *X*. Subject to the rate constraint *I*(*T*; *X*) = *C*, we seek to maximize *I*(*T*; *Y*). This constrained optimization problem can be written using a Lagrange multiplier as

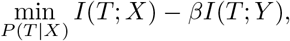

where the Lagrange multiplier *β* parametrizes the assumed rate *I*(*T*; *X*) = *C*.

When studying the predictive information bottleneck [41, 51, 56], the variables are the stimulus *S*_*t*_ as the input *X* and the future of the stimulus *S*_*t*+∆*t*_ as the relevance variable *Y*, giving the optimization problem

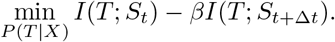

For our stochastic stimulus [48], the joint distribution *P* (*S*_*t*_, *S*_*t*+∆*t*_) is Gaussian, allowing us to solve the predictive information bottleneck problem exactly [15, 56] for arbitrary *β* and ∆*t*. Optimal encoders take the form of noisy linear measurements of the stimulus *T* = *AS*_*t*_ + *ξ*, where Σ_*ξ*_ = *I*_2_. In this Gaussian case, with the limitation of using only linear measurements of the stimulus state *S*_*t*_, it is also possible to exactly compute the privacy funnel [43], a lower bound on predictive information, which tells us how quickly information can be lost for some ∆*t*. To apply this approach to neural data, it is necessary to estimate the mutual information between the neural response and the stimulus both at equal times and in the future.

### Motion anticipation for IB compressions

The critically damped stochastic moving bar

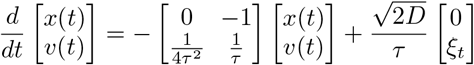

with time constant *τ* and diffusion constant 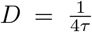, which fixes the position to have unit variance, evolves in a two-dimensional state-space *S*_*t*_ = (*x*_*t*_, *v*_*t*_) where each point in time is associated with a position and a velocity. For large amounts of compression, the IB encoding is a noisy linear map from the two-dimensional state-space down to one-dimension. These one-dimensional retinal encodings can be parametrized as *T*_*t*_(*θ, α*) = *α*[cos(*θ*), 2*τ* sin(*θ*)] · *S*_*t*_ + *ξ*_*t*_, where *ξ*_*t*_ is zero-mean, unit-variance Gaussian noise. The 2*τ* value appears in front of the velocity term to make the velocity variance 1. With this choice, 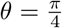is a compression that gives equal priority to position and velocity. The variable *α* sets a signal to noise ratio, and *θ* determines the relative importance of position and velocity. *θ* = 0 corresponds to a pure encoding of position, and *θ* = *π/*2 corresponds to a pure encoding of velocity. Taking *τ* = 1, and examining the zero noise limit, we have the following correlation functions:

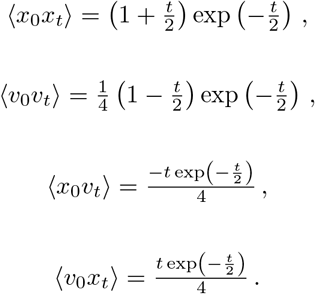

The correlation functions for the compression *T*_*t*_(*θ, α*) = *α*(*x*_*t*_ cos *θ* + 2*v*_*t*_ sin *θ*) + *ξ*_*t*_ are:

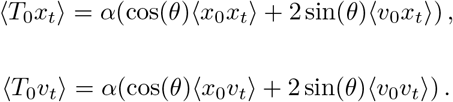

These can be used to calculate the fraction of position and velocity variance, 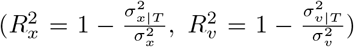, captured by the IB compression:

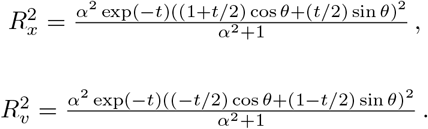

Differentiating 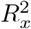 with respect to *t*, a critical point is found that corresponds to a maximum at

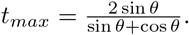

When *θ* is restricted to be between 0 (position-only encoding) and *π/*4 (position-and-velocity-equal encoding), the point in time that the compression’s correlation with position attains a maximum value that increases with *θ*, corresponding to increasing ∆*t*_IB_ in an IB problem.

### Stimulus mutual information estimation

We take avariational approach to the estimation of information. Considering the information between the stimulus *S* at time *t*, and the response *R* at time *t* with lag *k*, we have *I*(*S*_*t*_; *R*_*t*+*k*_) = *H*(*S*_*t*_)− *H*(*S*_*t*_|*R*_*t*+*k*_). The entropy of the stimulus can be computed exactly, as the steady-state distribution of the stimulus is Gaussian. Therefore,

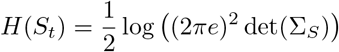

where Σ_*S*_ is the steady state-covariance for our stimulus. This same quantity can also be computed exactly for position *x*_*t*_ and velocity *v*_*t*_; as these are both Gaussian, we again have

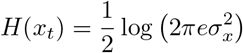

and

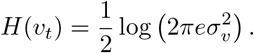

To estimate the mutual information we need only to estimate *H*(*S*_*t*_ *R*_*t*+*k*_).

This conditional entropy can be interpreted in terms of the performance of a decoder. Consider an estimate of the stimulus state 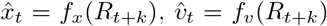, where reconstruction errors are given by 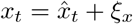 and 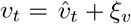 and *ξ*_*x*_ and *ξ*_*v*_ are additive noise terms that may differ in magnitude. Assuming Gaussianity and applying the law of total variance, we have for the conditional position variance

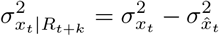

and the conditional velocity variance

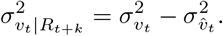

Applying the formula for the entropy of Gaussian variables to these quantities, we have

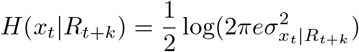

and

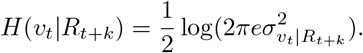

In the case that *f*_*x*_ and *f*_*v*_ are suboptimal decoders, the entropy will be overestimated, and so the estimate of mutual information will be a lower bound. If *f*_*x*_ and *f*_*v*_ are optimal decoders, then the entropy will be accurately estimated and the estimate of mutual information will be accurate. In general, how tight the variational lower bound on mutual information is depends on how powerful the inferred decoder is. Even though position and velocity are uncorrelated at equal times, the decoded estimates of position and velocity can be correlated. Therefore these correlations need to be taken into account when estimating the entropy *H*(*S*_*t*_). Correlations between position and velocity estimates are observed in IB optimal encodings.

Again, applying the law of total variance and assuming Gaussianity, we have

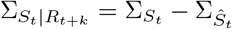

and

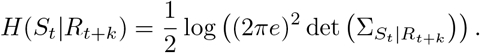

We note that when taking this approach to measuring information between stimulus and response it is important to measure mutual information for both the position and velocity. A measurement based only on position would understate the amount of information in the population. In addition, it can cause violations of the data processing inequality. For example, we observe that the peak in information between population response and the position occurs in the future for one stimulus condition, while the peak in total information remains in the past.

### Identifying the closest information bottleneck problem

The goal is to find the IB problem, defined by a prediction horizon ∆*t* and a compression level *β*, whose optimal encoding most closely matches what the retina actually does. We define ‘closest match’ as follows: we take the retina’s decoded posterior Σ_*S*|*R*_ at the lag which maximizes mutual information with the stimulus, and search over the family of IB-optimal posterior covariances 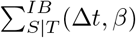 for the one that minimizes the KL-divergence with the empirical posterior subject to a rate-matching constraint.

Given the decoder estimate of the posterior covariance 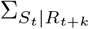, we treat *R*_*t*+*k*_ as the analog of the information bottleneck compression *T* and consider which predictive IB problem gives the most similar posterior distribution. Each predictive IB optimal encoder is parametrized by a prediction horizon ∆*t* associated with the relevance variable *S*_*t*+∆*t*_, and a tradeoff parameter value *β* that sets the allowed rate *I*(*T*; *S*_*t*_). These parameters define an IB optimal posterior covariance as a function of ∆*t* and *β* 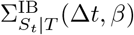. We define the closest IB problem as

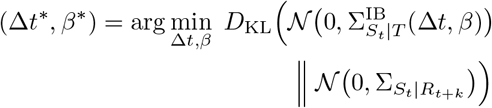

subject to the constraint

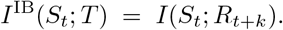

Note that

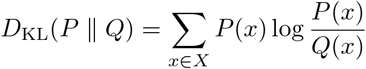

is asymmetric in its arguments. We treat the IB posterior as the reference distribution that is approximated by the empirical posterior, rather than the reverse. For that reason, we set 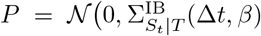 and 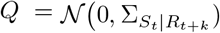.

Because both posteriors are zero-mean Gaussians, the KL divergence has a closed form in the two covariances alone. The rate-matching constraint fixes *β* as an implicit function *β*(∆*t*), reducing the search to a one-dimensional minimization over ∆*t*. Brent’s method [12] is used to minimize this function.

### Estimates of future stimulus information

The information bottleneck assumes the Markov relationship *T* −*X*− *Y*, implying that *H*(*Y*|*X*) ≤ *H*(*Y*|*T*). In our predictive information bottleneck problem, this means that in order to satisfy these assumptions we must have *H*(*S*_*t*+∆*t*_|*S*_*t*_) *H*≤ (*S*_*t*+∆*t*_|*R*_*t*+*k*_). The introduction of an offset lag *k*, similar to [41], can potentially create problems with this assumption. Our lag was selected to be the point in the future at which the retina population response has maximum information about the present stimulus state. While this choice is appropriate for the time at which a hypothetical downstream population of neurons could have the most information about the stimulus, it does not guarantee that *H*(*S*_*t*+∆*t*_|*S*_*t*_) ≤*H*(*S*_*t*+∆*t*_|*R*_*t*+*k*_). Therefore, in order to respect this assumption, we do not use a decoder based approach to estimate predictive mutual information. Rather, we work from our estimated 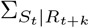 to estimate future information for arbitrary ∆*t*, in a way that, by construction, imposes the assumption *R*_*t*+*k*_ −*S*_*t*_ −*S*_*t*+∆*t*_. In addition to respecting the assumptions of the information bottleneck, this allows us to compute predictive information for arbitrary time ∆*t* rather than being limited to the temporal resolution of the stimulus and spiking response.

Decoding provides us with an estimate of the conditional covariance 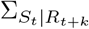. To get an estimate of 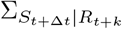, we begin by finding the regression coefficients with no compression. We know from the joint Gaussianity of the stimulus that

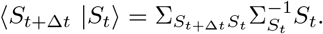

We call these coefficients for propagating the stimulus forward 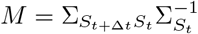. From the Schur complement, we can get the residual uncertainty about the future resulting from the stochasticity of the stimulus

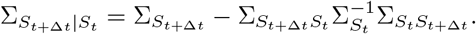

Now applying our Markov condition we also get the additional uncertainty resulting from compression

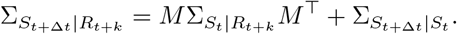

The conditional variances for position and velocity can be found by marginalizing the full conditional distribution. Estimates for the predictive mutual information can then be computed as

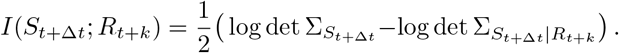

Similarly, for position and velocity alone, we have

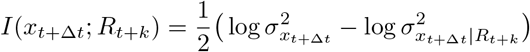

and

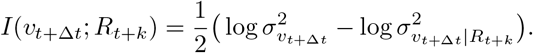

This approach differs from [41], where there is no guarantee that the compressions will respect the information bottleneck Markov assumption.

### Identifying the implied encoder

To draw comparisons between our inferred regression models and the solutions to the predictive information bottleneck problem, we use our conditional covariance matrix 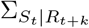 to find the matrix *A* that gives the same covariance matrix. We begin by noting that for the Gaussian information bottleneck encoder formula *T* = *AX* + *ξ*, where Σ_*ξ*_ = *I*_*x*_, the cross covariance matrix is

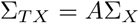

and the compression covariance is

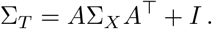

We substitute these into the equation for the conditional covariance

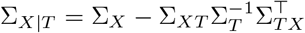

to get

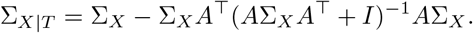

Applying the Woodbury matrix identity we note that

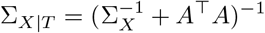

and therefore

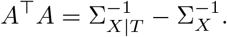

We can then get a canonical representation for our estimated encoding matrix *Â* by taking the eigenvectors *Q* and eigenvalues *λ* of 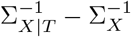 and setting

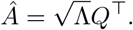

### Decoding position and velocity from the retina population response

Three different models were considered for decoding the stimulus position and velocity at time *t* from the population response at time *t* + *k*. As a baseline model we considered *L*_2_-regularized linear regression. Cross-validation was used to select regularization strength. To learn a set of linear features that could be used in more complex models, we used multi-target partial least squares regression to predict both position and velocity. Across multiple window sizes, performance saturated with 20 features.

For a more expressive, non-linear model, we use the features inferred from the partial least squares model to predict position and velocity with independent Gaussian processes per target. To handle the large number of training samples, we adopt a variational approach with 256 inducing points initialized from a random subset of the training data [31, 66], as implemented in GPyTorch [25]. Inputs and targets are standardized prior to training, and predictions are projected back to the original space. We use a constant mean and a squared-exponential covariance kernel with automatic relevance determination.

As the Gaussian Process model offered a significant performance improvement over the linear models in decoding both position (Fig. S1) and velocity (Fig. S2) we use it as the decoding model discussed in the main text. In addition to showing lower overall performance measured by *R*^2^ for position and velocity, the linear models also show evidence of a specified mean for both position (Fig. S3) and velocity (Fig. S4). All models show evidence for homoskedastic errors in both position (Fig. S3) and velocity (Fig. S4). The accuracy of the mutual information estimates depends on the Gaussianity of the residual distribution. To evaluate this we show QQ plots for both position and velocity across all models (Fig. S5), along with estimated values for excess kurtosis and skewness. QQ plots compare quantiles of the empirical residual distribution to those of a Gaussian distribution. In all models and stimulus conditions the Gaussian assumption for the residuals is reasonable (Fig. S6). Additionally, a decoder-based estimate of mutual information (training a new decoder for every future time considered) is compared to the analytic uncertainty approach discussed above. The analytic uncertainty approach is more conservative, and respects the information bottleneck assumptions by construction (Fig. S7).

All models are trained using five-fold cross validation, and a different model is trained for each stimulus condition and for each choice of relative stimulus-response lag *k*. To ensure that test data is not contaminated by temporal correlations, data is shuffled by trial. The training sets used consist of 24 unique trials, and each test set consists of 6 unique trials. Quantities derived from decoders are shown as averages over folds, with standard deviations also taken over folds.

### Entropy estimation

To estimate the entropy of the neural population, we use the cross-entropy estimate found from inferred Restricted Boltzmann Machines. This gives an upper bound on the true entropy [45]. These expressive models have been found to accurately describe retinal population activity [24, 69]. The empirical cross-entropy is an unbiased estimator of the true cross-entropy and bounds the true entropy from above. The gap between the cross-entropy and the population entropy is the Kullback-Leibler divergence between the underlying distribution and the model, which is expected to be small for such an expressive model. We use five-fold cross-validation over stimulus trials and report the average entropy on held-out test data as the estimate; the reported error bars are the standard deviations over folds. Folds are shuffled over trial indices instead of all data to manage contamination from temporal correlation.

As Restricted Boltzmann Machines are unnormalized statistical models, we need to estimate the partition function to report the cross-entropy. We use the Good-Turing estimator to estimate the partition function [28], as discussed in [30]. For optimization, we use the minimum probability flow approach [62], and use model samples with the factorization assumption to construct a connectivity matrix [61].

We used the cross-validation performance to select the appropriate number of hidden units. Model convergence was additionally assessed by comparing model covariance to data covariance, and model spike count distributions to data spike count distributions. All reported entropy measures based on these models are reported from the average of held-out test data. The error bars are the standard deviations over folds.

### Stimulus and population surprise

For the stimulus surprise we know that the steady-state distribution is 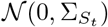, therefore we can straightforwardly compute the quantity ™ log *p* exactly as

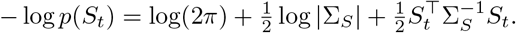

The compressed representation is the linear encoding *T* = *AS*_*t*_ + *ξ* with *ξ* ∽ 𝒩 (0, *I*), so marginally *T* ∽ 𝒩 (0, Σ_*T*_) with Σ_*T*_ = *A*Σ_*S*_*A*^⊤^+ *I*. Because *T* is stochastic, we report the expected surprise averaged over the encoding noise

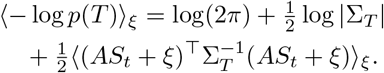

Expanding the quadratic and using ⟨*ξ*⟩ = 0, ⟨*ξξ*^⊤^⟩ = *I*,

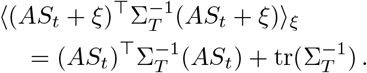

Therefore,

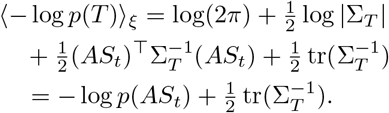

The noise contributes only the constant offset 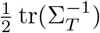,so all time variation comes from the noiseless encoding *AS*_*t*_.

For the retina population response, the negative unnormalized log probability, or energy, was used because the additive constant from the partition function is irrelevant for this analysis. For the comparison between the population response surprise and the compressed and uncompressed stimulus surprise, a gain and an offset are used for plotting, but this does not affect the correlation between the quantities. For example traces of all surprise comparisons see (Fig. S8).

### Retina reversal response

Previously the retina reversal response has been probed with deterministic stimuli [60]. To develop a probabilistic notion of a reversal response we consider as a function of the present stimulus state *S*_*t*_ = (*x*_*t*_, *v*_*t*_) the probability that the velocity will change sign ∆*t*_rev_ in the future. We introduce the random variable Rev_∆*t*_ that is 1 when sign(*v*_*t*_) ≠ sign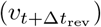 and 0 otherwise.

To compute the probability of a reversal *P* (Rev_∆*t*_ = 1) we have a version of the orthant problem [26] for a 2D Gaussian. Applying Sheppard’s formula [26], we know that

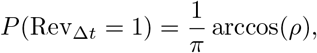

where *ρ* is the correlation in the joint distribution 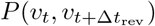. For the reversal probability conditioned on the stimulus, the probability only depends on the CDF Φ of the conditional distribution 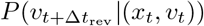 and we have

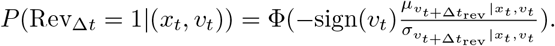

The entropy of the unconditional distribution *P* (Rev_∆*t*_) can be computed exactly, and the entropy of the conditional distribution *P* (Rev_∆*t*_ |(*x*_*t*_, *v*_*t*_)) can be computed with numerical integration, therefore we can estimate the information between the stimulus and Rev_∆*t*_.

To compute the same quantity for the compressed stimulus we consider a noiseless 1D projection *T*_*θ*_ = (cos *θ*)*x*_*t*_ + (2*τ* sin *θ*)*v*_*t*_. The 2*τ* factor puts us into whitened coordinates so *θ* = 0 is position encoding and 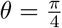 corresponds to equal encoding of position and velocity. We can compute *P* (Rev_∆*t*_ = 1 |*T*_*θ*_), by breaking the problem down into two orthant problems

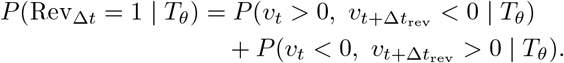

In turn each of these problems can be broken down as follows

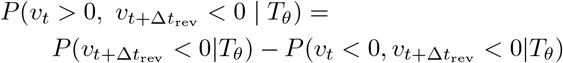

and

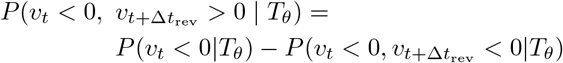

therefore

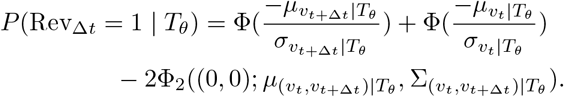

where Φ_2_ is the CDF for a 2D Gaussian with mean 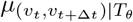 and covariance matrix 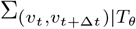. And so, via numerical integration, we can also compute the information between the compressed stimulus and Rev_∆*t*_.

For our stochastic moving-bar stimulus with IB relevance variable *S*_*t*+∆*t*_, the IB encoding incorporates monotonically more velocity information, up to equal amounts of position and velocity as ∆*t* goes to infinity [56]. Sweeping the projection angle *θ* from 0 to *π/*4 shows that, across a range of ∆*t*_rev_, the compressions appropriate for larger IB relevance horizons carry more information about Rev_∆*t*_ (Fig. S9).

## Acknowledgments

KB would like to thank Tobias Kühn for useful discussions, and Sophie Colt for helpful comments on the manuscript. This work was supported by the Physics Frontier Center for Living Systems through the National Science Foundation award NSF PHY-2317138; the NSF-Simons National Institute for Theory and Mathematics in Biology, awards NSF DMS-2235451 and Simons Foundation MP-TMPS-00005320; the University of Chicago Materials Research Science and Engineering Center, award NSF DMR-2011854; the Center for the Physics of Biological Function, NSF PHY-1734030 and by the Polymaths Program from Schmidt Sciences, LLC, to SEP.

---

**FIG. S1.**
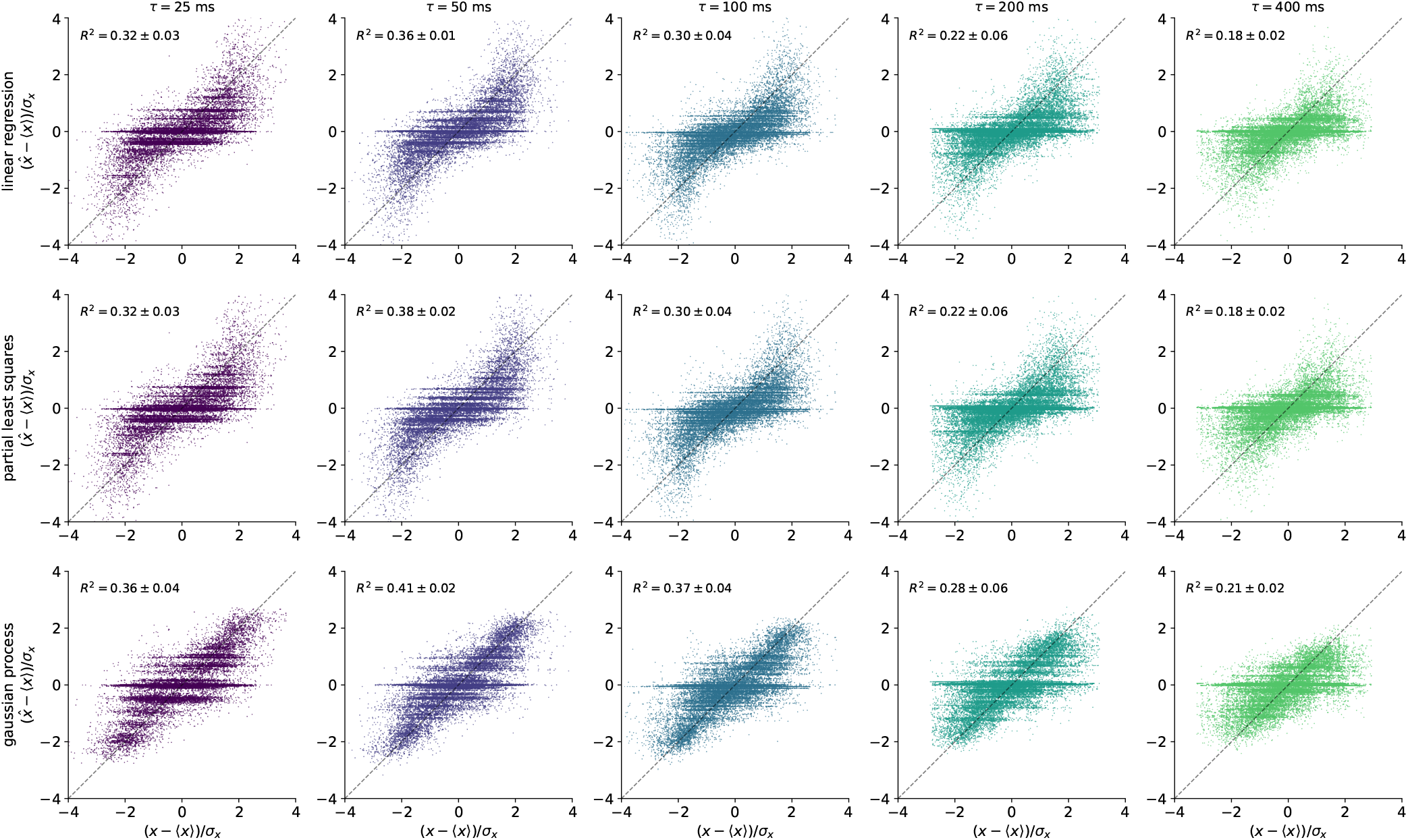
Decoder performance for position across models and stimuli conditions. True normalized position values are compared to predicted position values from the linear regression model (top row), the partial least squares model (middle row) and the gaussian process model (bottom row). Model shown is for choice of relative stimulus response lag that maximizes total (position and velocity) mutual information.

**FIG. S2.**
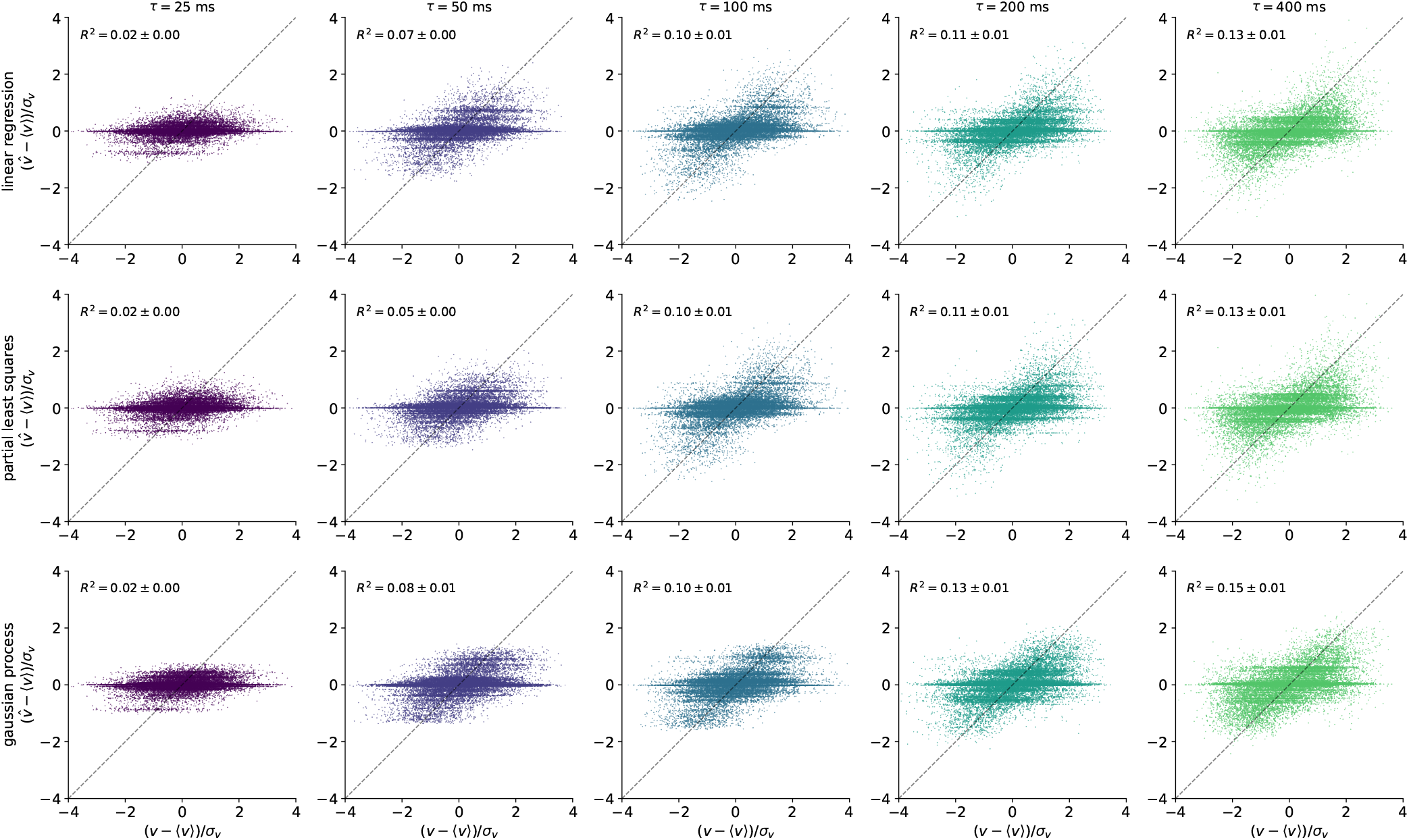
Decoder performance for velocity across models and stimuli conditions. True normalized velocity values are compared to predicted velocity values from the linear regression model (top row), the partial least squares model (middle row) and the gaussian process model (bottom row). Model shown is for choice of relative stimulus response lag that maximizes total (position and velocity) mutual information.

**FIG. S3.**
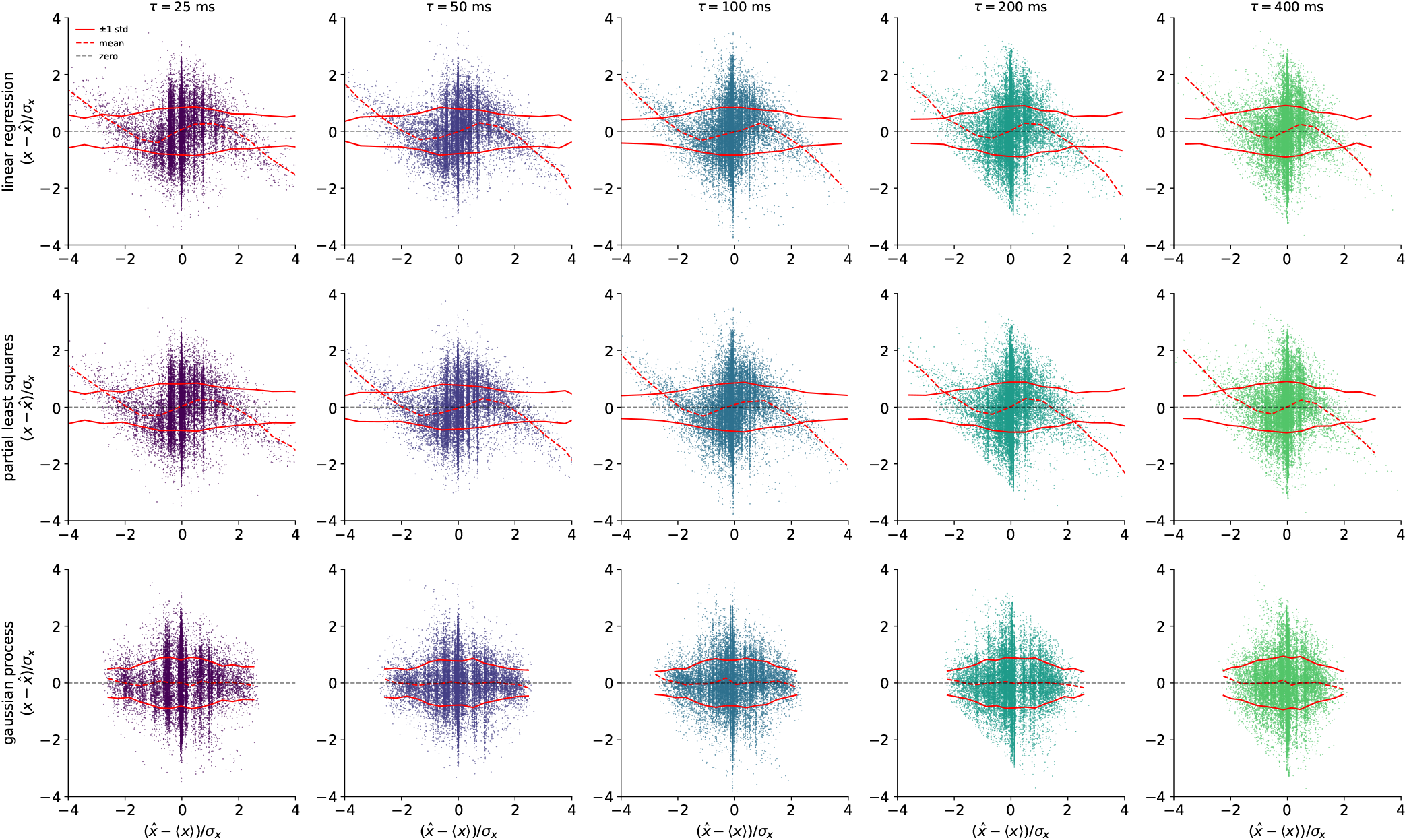
Position residuals across models and stimuli conditions. Normalized position residuals are shown as a function of normalized position value for the linear regression model (top row), the partial least squares model (middle row) and the gaussian process model (bottom row). Model shown is for choice of relative stimulus response lag that maximizes total (position and velocity) mutual information. Dashed red line indicates mean of residuals as a function of normalized position. Solid red line indicates residual standard deviation as a function of normalized position. Means and standard deviations were computed for fifteen linearly spaced bins.

**FIG. S4.**
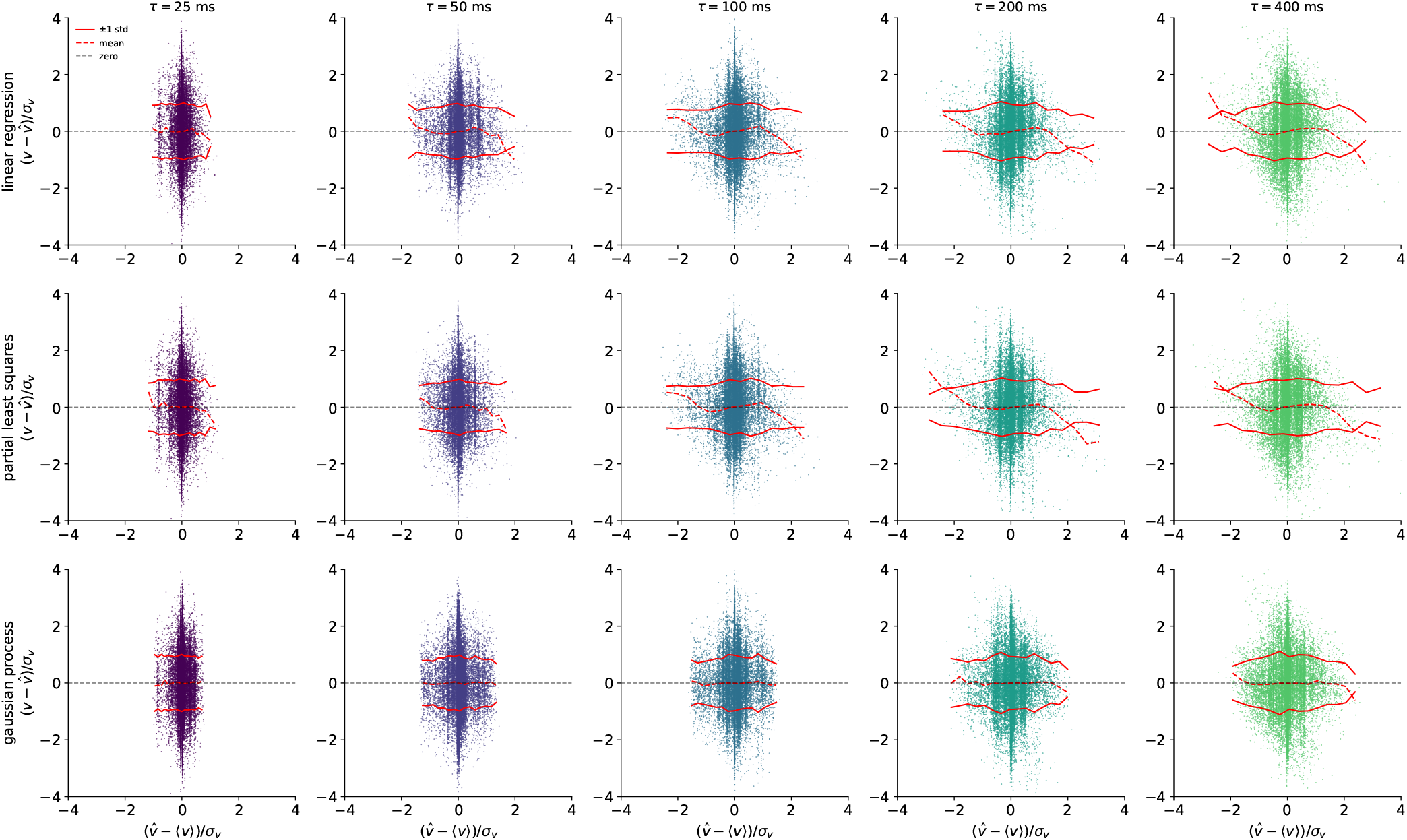
Velocity residuals across models and stimuli conditions. Normalized velocity residuals are shown as a function of normalized velocity value for the linear regression model (top row), the partial least squares model (middle row) and the gaussian process model (bottom row). Model shown is for choice of relative stimulus response lag that maximizes total (position and velocity) mutual information. Dashed red line indicates mean of residuals as a function of normalized velocity. Solid red line indicates residual standard deviation as a function of normalized velocity. Means and standard deviations were computed for fifteen linearly spaced bins.

**FIG. S5.**
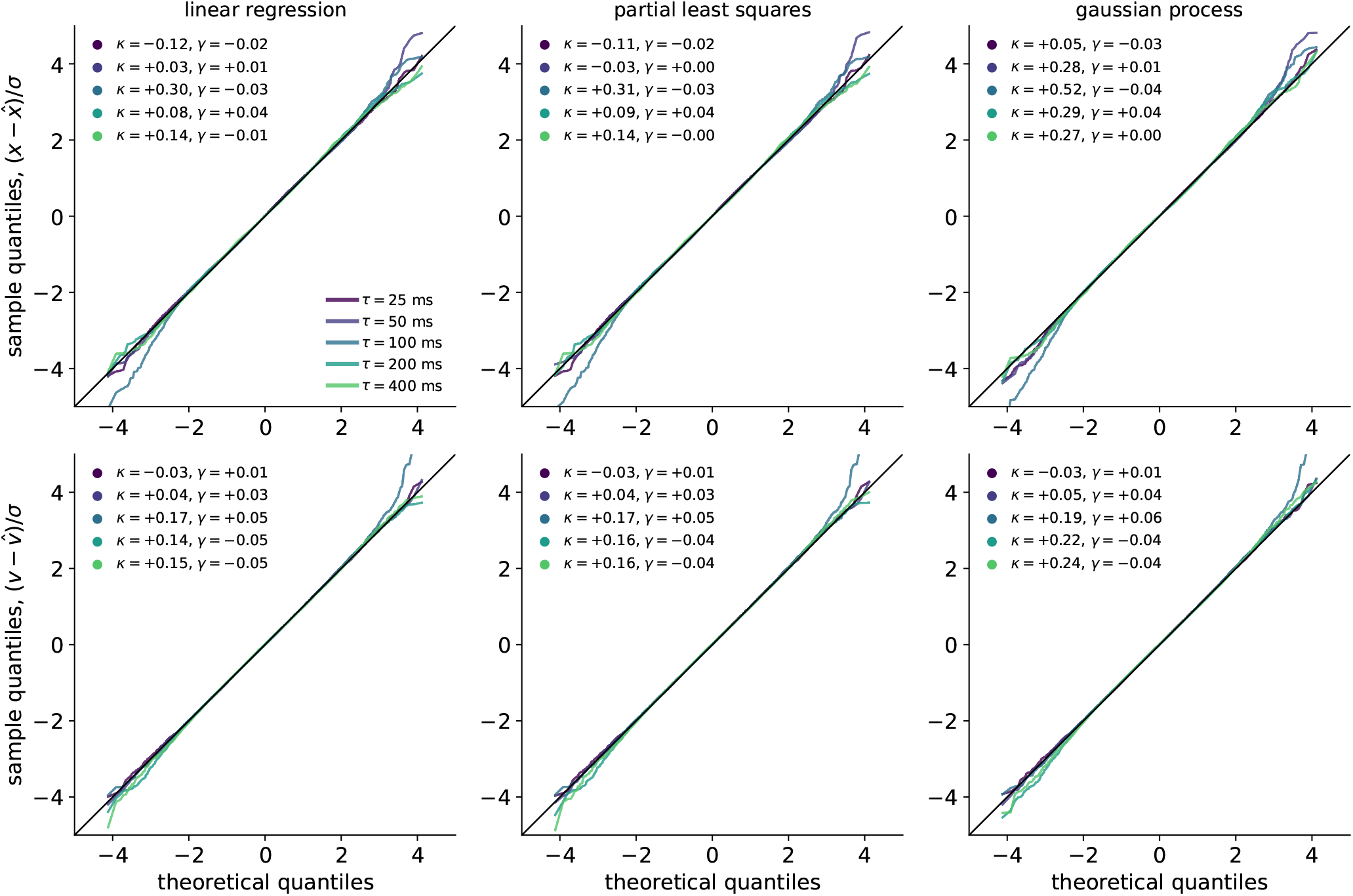
Gaussianity of residuals for position and velocity. QQ plots comparing normalized residual quantiles to corresponding Gaussian quantiles across all stimulus conditions and decoder models. Excess kurtosis and skewness are also shown for each decoder and stimulus condition.

**FIG. S6.**
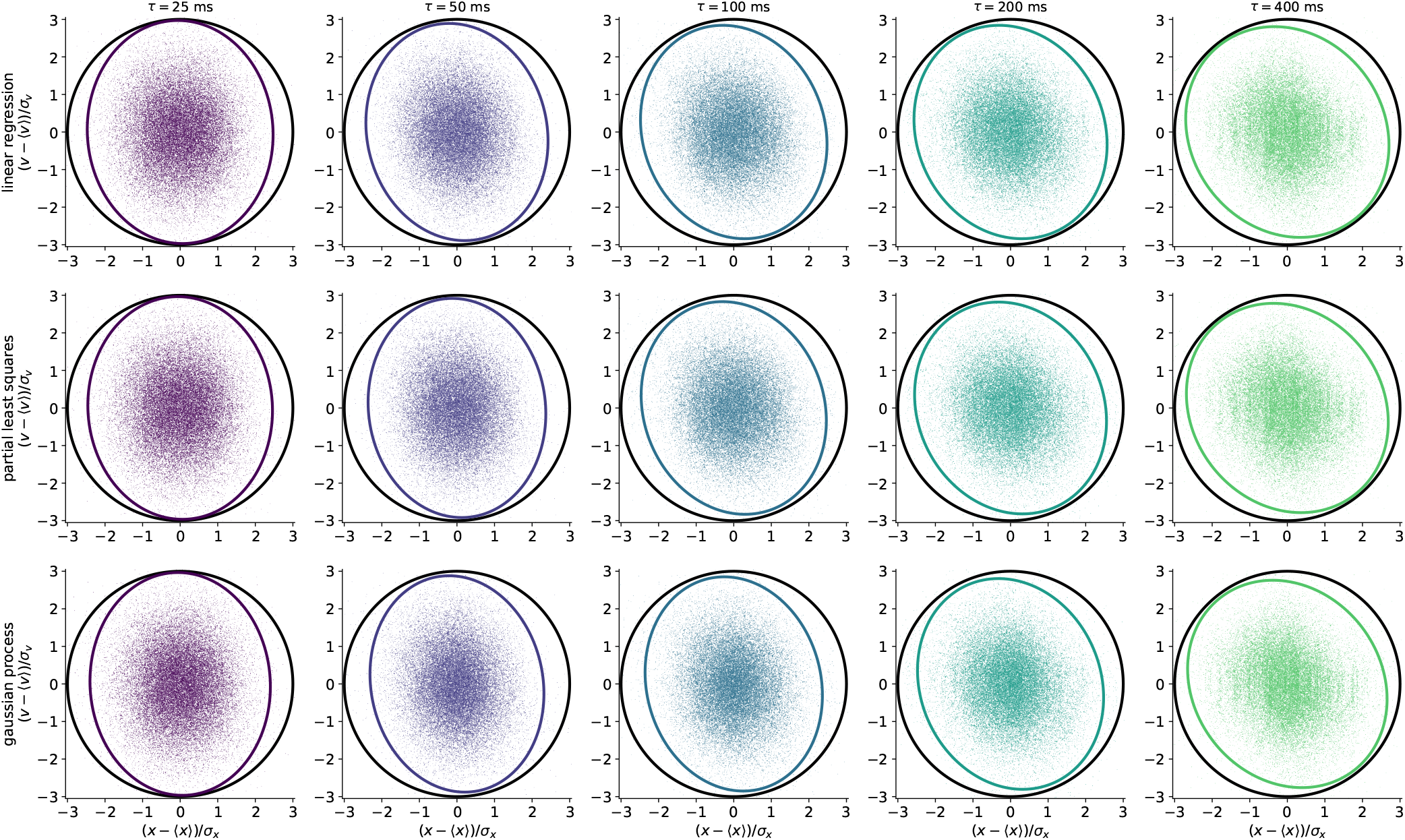
Error distribution for all models and stimuli conditions. A scatter plot shows the distribution of normalized decoding errors across stimuli conditions and decoding models. The black ellipse shows the three standard deviation contour for an uninformative decoder, or the normalized steady-state variance 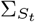. The colored ellipse show the expected three standard deviation contour for the different decoders, which has covariance 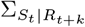. In all cases model shown is for choice of relative stimulus response lag that maximizes total (position and velocity) mutual information.

**FIG. S7.**
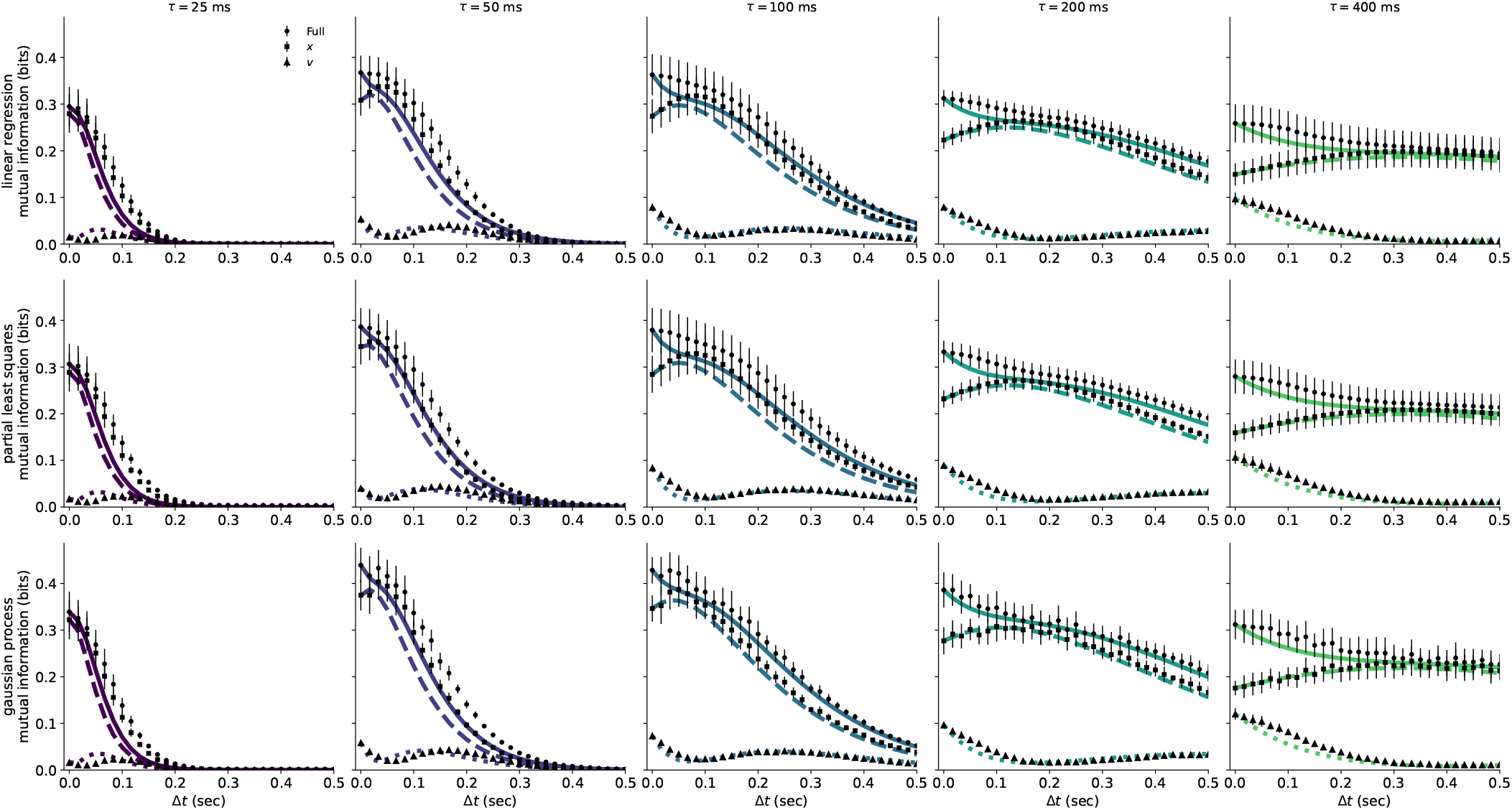
Future mutual information estimation: analytic uncertainty vs decoder. Position, velocity, and total mutual information as a function of ∆*t* for the described analytic uncertainty method (lines) and the decoder approach (points). Results are shown for all models and stimulus conditions. Error bars on the future decoder method are standard deviations over folds.

**FIG. S8.**
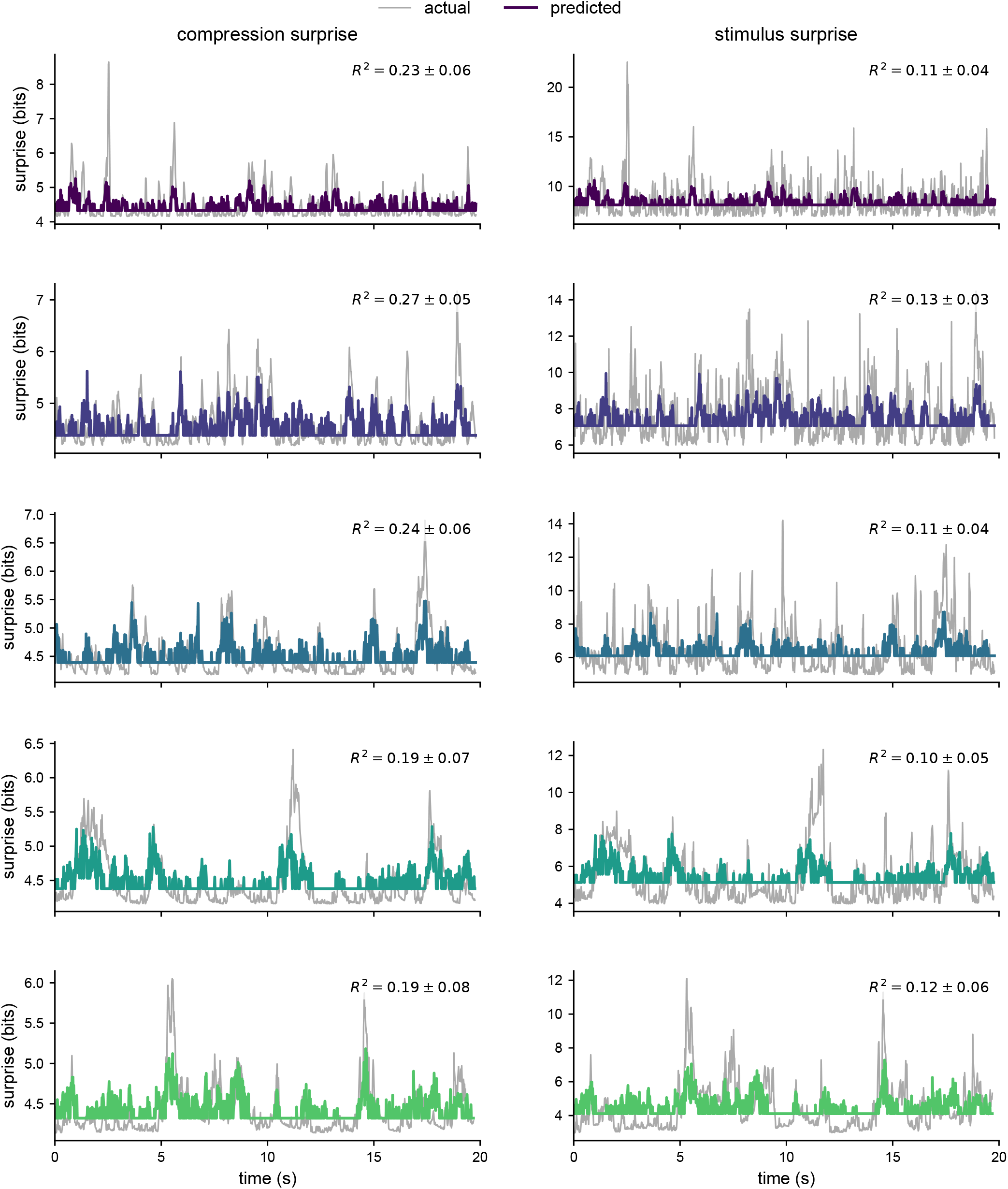
Surprise comparison: Full comparisons are shown for the population response surprise and the compressed and uncompressed stimulus surprise, similar to (Fig. 5) b) for a full trial from each stimulus condition. *R*^2^ values are again averages over all trials with reported errors taken as the standard deviation over trials.

**FIG. S9.**
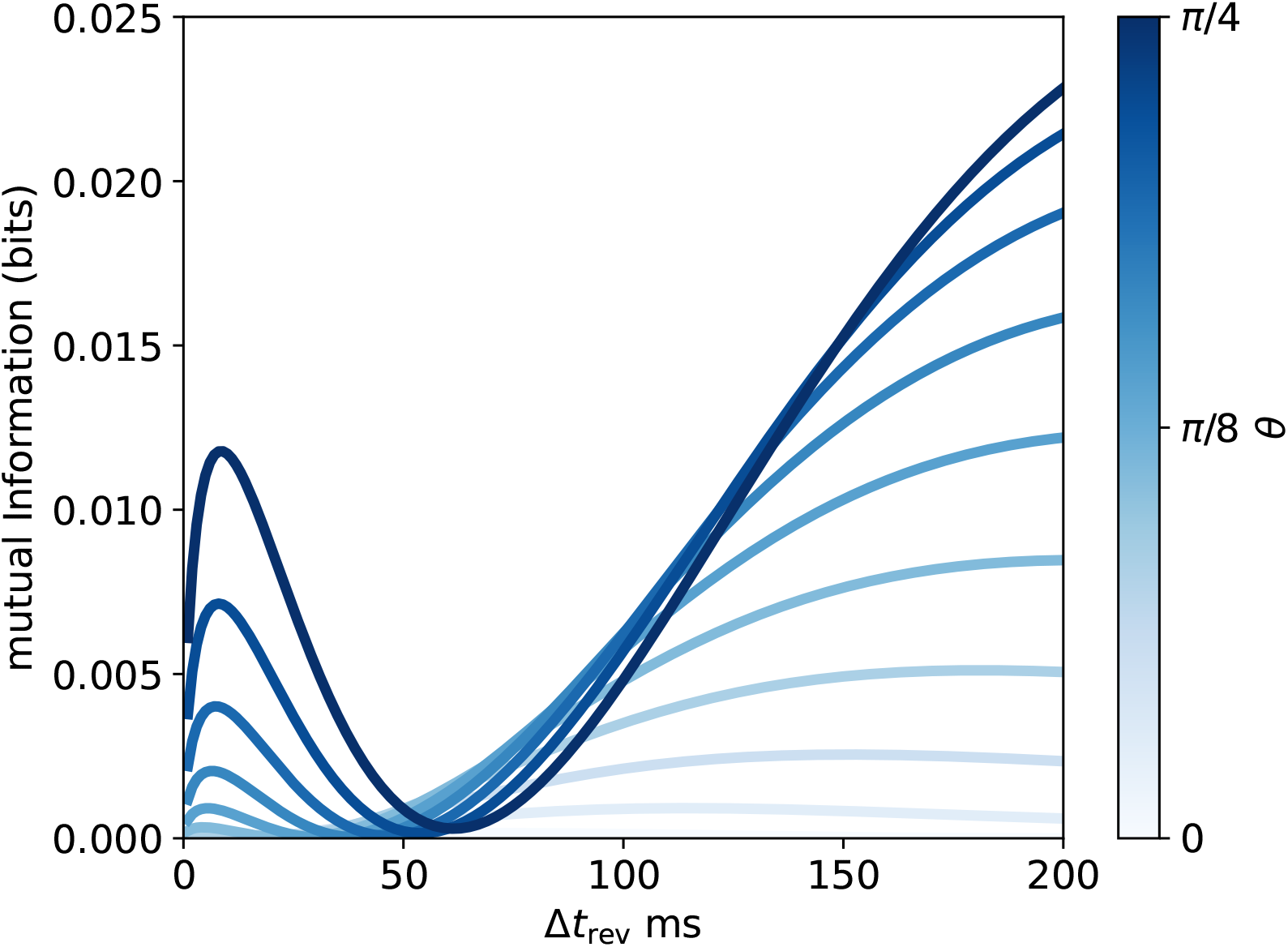
Reversal information: Mutual information between compressed stimulus and reversal events as a function of reversal horizon ∆*t*_rev_ for multiple different compression schemes. Example is for *τ* = 100 ms.

## Notes

### Competing Interest Statement

The authors have declared no competing interest.

### Summary of Updates

Methods expanded and figures updated in response to requested revisions.

